# The Allen Cell and Structure Segmenter: a new open source toolkit for segmenting 3D intracellular structures in fluorescence microscopy images

**DOI:** 10.1101/491035

**Authors:** Jianxu Chen, Liya Ding, Matheus P. Viana, HyeonWoo Lee, M. Filip Sluezwski, Benjamin Morris, Melissa C. Hendershott, Ruian Yang, Irina A. Mueller, Susanne M. Rafelski

## Abstract

A continuing challenge in quantitative cell biology is the accurate and robust 3D segmentation of structures of interest from fluorescence microscopy images in an automated, reproducible, and widely accessible manner for subsequent interpretable data analysis. We describe the Allen Cell and Structure Segmenter (Segmenter), a Python-based open source toolkit developed for 3D segmentation of cells and intracellular structures in fluorescence microscope images. This toolkit brings together classic image segmentation and iterative deep learning workflows first to generate initial high-quality 3D intracellular structure segmentations and then to easily curate these results to generate the ground truths for building robust and accurate deep learning models. The toolkit takes advantage of the high-replicate 3D live cell image data collected at the Allen Institute for Cell Science of over 30 endogenous fluorescently tagged human induced pluripotent stem cell (hiPSC) lines. Each cell line represents a different intracellular structure with one or more distinct localization patterns within undifferentiated hiPS cells and hiPSC-derived cardiomyocytes. The Segmenter consists of two complementary elements, a classic image segmentation workflow with a restricted set of algorithms and parameters and an iterative deep learning segmentation workflow. We created a collection of 20 classic image segmentation workflows based on 20 distinct and representative intracellular structure localization patterns as a “lookup table” reference and starting point for users. The iterative deep learning workflow can take over when the classic segmentation workflow is insufficient. Two straightforward “human-in-the-loop” curation strategies convert a set of classic image segmentation workflow results into a set of 3D ground truth images for iterative model training without the need for manual painting in 3D. The deep learning model architectures used in this toolkit were designed and tested specifically for 3D fluorescence microscope images and implemented as readable scripts. The Segmenter thus leverages state of the art computer vision algorithms in an accessible way to facilitate their application by the experimental biology researcher.

We include two useful applications to demonstrate how we used the classic image segmentation and iterative deep learning workflows to solve more challenging 3D segmentation tasks. First, we introduce the ‘Training Assay’ approach, a new experimental-computational co-design concept to generate more biologically accurate segmentation ground truths. We combined the iterative deep learning workflow with three Training Assays to develop a robust, scalable cell and nuclear instance segmentation algorithm, which could achieve accurate target segmentation for over 98% of individual cells and over 80% of entire fields of view. Second, we demonstrate how to extend the lamin B1 segmentation model built from the iterative deep learning workflow to obtain more biologically accurate lamin B1 segmentation by utilizing multi-channel inputs and combining multiple ML models. The steps and workflows used to develop these algorithms are generalizable to other similar segmentation challenges. More information, including tutorials and code repositories, are available at allencell.org/segmenter.

## Introduction

Modern fluorescence microscopy has revolutionized imaging of tissues, cells, subcellular structures, and proteins [Kervrann et al., 2016]. The resulting multi-dimensional image data (3D, time-lapse, multiple imaging channels, or combinations thereof, etc.) require further analysis with a variety of qualitative and quantitative approaches. Simple visual inspection of small image data sets is used to rapidly assess general image quality or compare differences among experimental conditions. Quantitative and automated analysis approaches, however, become necessary when the number of images is large, the differences between experimental conditions are subtle or complex, or the image data and their interpretations are used to develop data-driven analyses and models. Quantifying images becomes especially important when dealing with 3D image data where even a straightforward comparison between two conditions can be difficult without quantitative measurements. Segmentation, the identification of every pixel (or voxel) that is either part or not part of that object, is key to extracting interpretable, quantitative measurements of an object in an image, permitting measurement of size, shape, number of objects and intensity of a given object, for example.

The large number of different sizes and shapes of structures found in cells makes image segmentation particularly challenging. Furthermore, 3D image data are inherently harder to work with than 2D images, presenting an additional challenge for cellular image segmentation and analysis. Existing 3D image segmentation methods can be categorized as classic image processing algorithms, traditional machine learning, and deep learning methods. Classic image processing algorithms are the most widely used by the cell biological research community and are accessible in two main ways. Some algorithms are available as collections of basic functions in several open platforms, including the widely used ImageJ [Schindelin et al., 2012], CellProfiler [Carpenter et al., 2006, McQuin et al., 2018], Icy [De Chaumont et al., 2012], and ITK-SNAP [Yushkevich et al., 2006]. However, basic functions in the open platforms are often not sufficiently accurate or up to date [Jerman et al., 2016]. Other published algorithms may be designed for a specific structure in a specific imaging modality and are often implemented and released individually [Neila et al., 2016, Hodneland et al., 2013, Smith and Barton, 2014]. Compared to general image processing platforms, such tools are less broadly applicable and often less convenient to apply.

Machine learning algorithms can also facilitate segmentation of 2D and 3D fluorescence microscopy images. Traditional machine learning algorithms, e.g., random forest and supporting vector machine, have been integrated successfully into openly accessible tools such as trainable WEKA segmentation [Arganda-Carreras et al., 2017] in ImageJ and ilastik [Sommer et al., 2011]. Users simply manually paint on selective pixels/voxels as foreground and background samples to create a ground truth training set. A traditional machine learning model is then automatically trained and applied on all selected images. While easy to use, these traditional machine learning models and tools are less effective than deep learning models especially when segmenting objects with occluded boundaries (e.g., tightly packed or highly textured objects [Çiçek et al., 2016, Chen et al., 2016B, Chen et al., 2017]. Unfortunately, deep learning models are “training data hungry.” Thus, tedious manual painting in 3D quickly becomes prohibitive for generating sufficient 3D ground truth data. Additionally, even if an adequate 3D ground truth can be prepared, access to convenient tools for building/deploying these deep learning models is currently prohibitive for many biologists. Existing tools, such as NiftyNet [Gibson et al., 2018] or DLTK [Pawlowski et al., 2017], are difficult to use without sufficient experience in deep learning and computer vision. Other tools, e.g., Aivia Cloud [DRVISION, 2018], are easier to use but not openly accessible.

The Allen Institute for Cell Science has developed a pipeline that generates high-replicate, dynamic image data on cell organization and activities using a collection of endogenous fluorescently tagged human induced pluripotent stem cell (hiPSC) lines (Allen Cell Collection; allencell.org; [Roberts et al., 2017, Viana et al., 2020]). Most lines express a monoallelic mEGFP-tagged protein that represents a particular intracellular structure (exceptions are the tagged sialyltransferase 1 line and the Ras-related protein Rab5-a line, which are also available as biallelic lines, and the centrin-2 and the CAAX domain of K-RAS (CAAX) lines, which are tagged with mTagRFP-T). To enable quantitative image and data analyses, we generated accurate and robust segmentations for over 30 intracellular 3D structure localization patterns. By developing and testing a diverse set of traditional segmentation algorithms on a wide range of intracellular structures, we identified a conceptually simple “classic image processing workflow” involving a limited number of classic image processing steps and algorithms that generated high-quality segmentations of these structures. These segmentations permitted initial analyses of basic morphometric features of these structures including size, number, shape, and location within the cell, and form the basis for more complicated feature parameterizations (**Fig. 1A**). To enhance the accuracy and robustness of these segmentations, we also developed a novel “iterative deep learning workflow” (**Fig. 1B**) that takes advantage of these high-quality classic segmentation results and applies them as an initial ground truth in an iterative deep learning-based approach to image segmentation.

**Figure 1:**
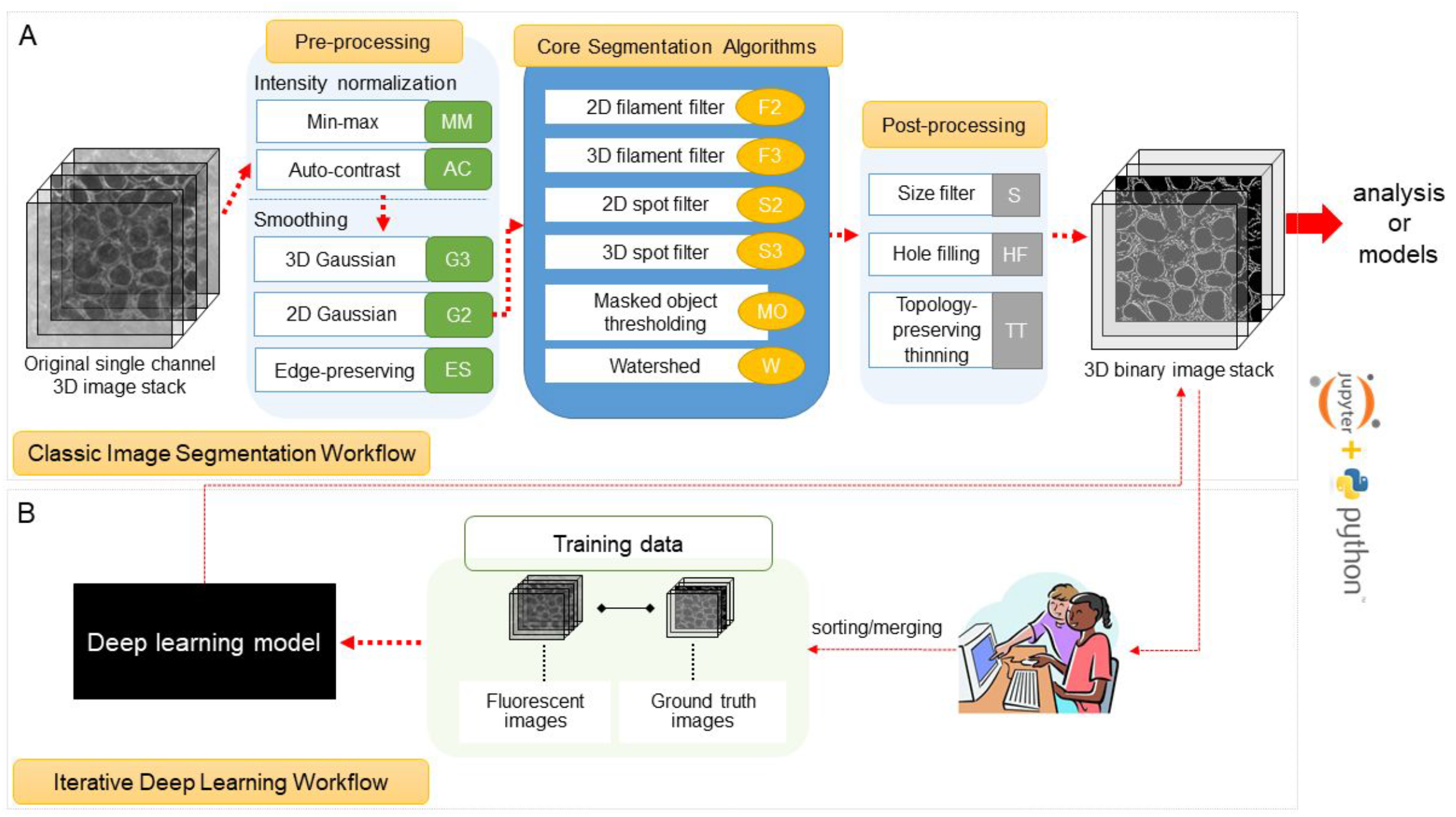
Overview of the Allen Cell and Structure Segmenter. **(A)** The classic image segmentation workflow consists of three steps and includes a restricted set of image processing algorithm choices and tunable parameters. The number of choices may grow as we develop more workflows. The updates will be available on allencell.org/segmenter **(B)** The iterative deep learning workflow is used when the accuracy or robustness of the classic image segmentation workflow is insufficient. Two human-in-the-loop strategies, sorting and merging, can be iteratively applied to build 3D ground truth training sets from the classic image segmentation workflow results or resultant deep learning 3D segmentation models.

These workflows are packaged into the Allen Cell and Structure Segmenter (Segmenter), an open-source, Python-based toolkit for segmentation of 3D microscopy images, building upon Numpy [Harris et al., 2020], Scipy [Virtanen et al., 2020], Scikit-image [Van de Walt et al., 2014], and ITK [McCormick et al., 2014] with many customizations. We developed this toolkit to make both workflows accessible to cell biologists wishing to quantify and analyze their own image data. The Segmenter offers two key advantages over other image processing packages. First, the classic image segmentation workflow streamlines algorithm and parameter choice. Users are provided with a “lookup table” of classic image segmentation workflows for 20 intracellular structure localization patterns with varying morphological properties that can be used as a starting point for segmentation. Second, the iterative deep learning workflow provides users with tools to apply results from the classic segmentation workflow to generate ground truth segmentations for training deep learning models, without manual painting, and then to use these models to iteratively improve those segmentation results.

We further extended the iterative deep learning workflow via a co-design of computational algorithms and biological experiments towards creating more biologically correct and robust segmentations (based on a segmentation target). We call this co-design the ‘Training Assay’ approach. The iterative deep learning workflow requires an accurate preliminary segmentation result to start with. However, often a biological image-based assay may be constrained by imaging requirements of the experimental system (such as requirement for reduced light in live cell imaging). Thus, the resultant assay images may be difficult to use directly to create these accurate preliminary segmentations. The iterative deep learning workflow permits a secondary assay, more amenable to accurate segmentation, to be linked to the primary image-based assay. In this way a segmentation model can be trained that achieves the better accuracy and robustness possible with the images of the secondary assay even when run on the poorer-quality primary assay images. For instance, it is difficult to accurately segment the 3D boundaries of DNA dye-labeled interphase nuclei in tightly packed, epithelial-like, hiPSC colonies, especially in the z direction, due to limitations in microscopy optics including the diffraction of light. However, the nuclear envelope visualized via lamin B1 creates a “shell” around the nucleus and thus permits a more biologically accurate detection of the nuclear boundary, especially in z. In this way we can image both the lamin B1 and the DNA dye channels for the same nuclei and use the more accurate segmentations obtained from the lamin B1 images as the ground truth segmentations to train a deep learning model to segment the nuclei via the DNA dye channel. This way we could now generate a more accurate and robust segmentation model to apply to other DNA-dye labeled images even in the absence of tagged lamin B1 in those cells.

We combined the iterative deep learning workflow with three Training Assays to develop a robust, scalable cell and nuclear instance segmentation algorithm for high-resolution 3D spinning disk confocal images of live hiPSC colonies from cell lines in the Allen Cell Collection. These cells were all labeled with a DNA dye (NucBlue Live) and a cell membrane dye (Cell Mask Deep Red). We achieved accurate target segmentations for over 98% of individual cells and over 80% of entire fields of view (FOVs; about 12 cells per image). This combination of iterative deep learning with the Training Assay approach has now permitted us to segment all cells and nuclei in our 3D microscopy images, a fundamental challenge to performing image-based single cell analysis at scale. The cell and nuclear segmentation on 18,186 FOVs have been released via the hiPSC Single-Cell Image Dataset [Viana et al., 2020] (https://open.quiltdata.com/b/allencell/packages/aics/hipsc_single_cell_image_dataset). This general approach and the steps outlined for this specific 3D cell and nuclear segmentation algorithm should generalize to other cell types and microscope image modalities.

Moreover, we present a method that builds onto a lamin B1 segmentation model trained from iterative deep leaning workflows to create an improved lamin B1 segmentation algorithm, by taking advantage of multi-channel input images and employing multiple complementary deep learning segmentation models. The improved lamin B1 segmentation algorithm is designed to obtain more biologically accurate segmentation results. This demonstrated the potential of Segmenter in solving challenging 3D segmentation problems to facilitate downstream quantitative biological analyses.

## Results and Discussion

### General overview of the Allen Cell and Structure Segmenter

The goal of the Allen Cell and Structure Segmenter (Segmenter) is to make flexible, robust, state-of-the-art 3D segmentation methods accessible to cell biology researchers. While developed specifically for the 3D segmentation of intracellular structures, the Segmenter may also be applicable to a variety of other image segmentation applications. It seamlessly integrates a traditional image segmentation workflow and an iterative deep learning workflow to streamline the segmentation process (**Fig.1**). The classic image segmentation workflow is based on a restricted set of both standard and cutting-edge algorithms and tunable parameters (**Fig.1A**) that we identified to be optimal for segmenting over 30 different intracellular structure localization patterns. We created a suite of 20 intracellular structure segmentation workflows which we present in a lookup table as a starting point for users to solve their own segmentation tasks (**Fig. 2**). The set of algorithms and the lookup table will continue to grow as we develop segmentation algorithms for more intracellular structures. New updates will be released on allencell.org/segmenter. In the iterative deep learning workflow of the Segmenter, we describe two new strategies for preparing 3D ground truth images without laborious and subjective manual painting in 3D (**Fig. 1B**). The training and testing of the deep learning model are customized for intracellular structures in 3D fluorescence microscopy images and implemented as readable scripts for researchers without experience in deep learning.

**Figure 2:**
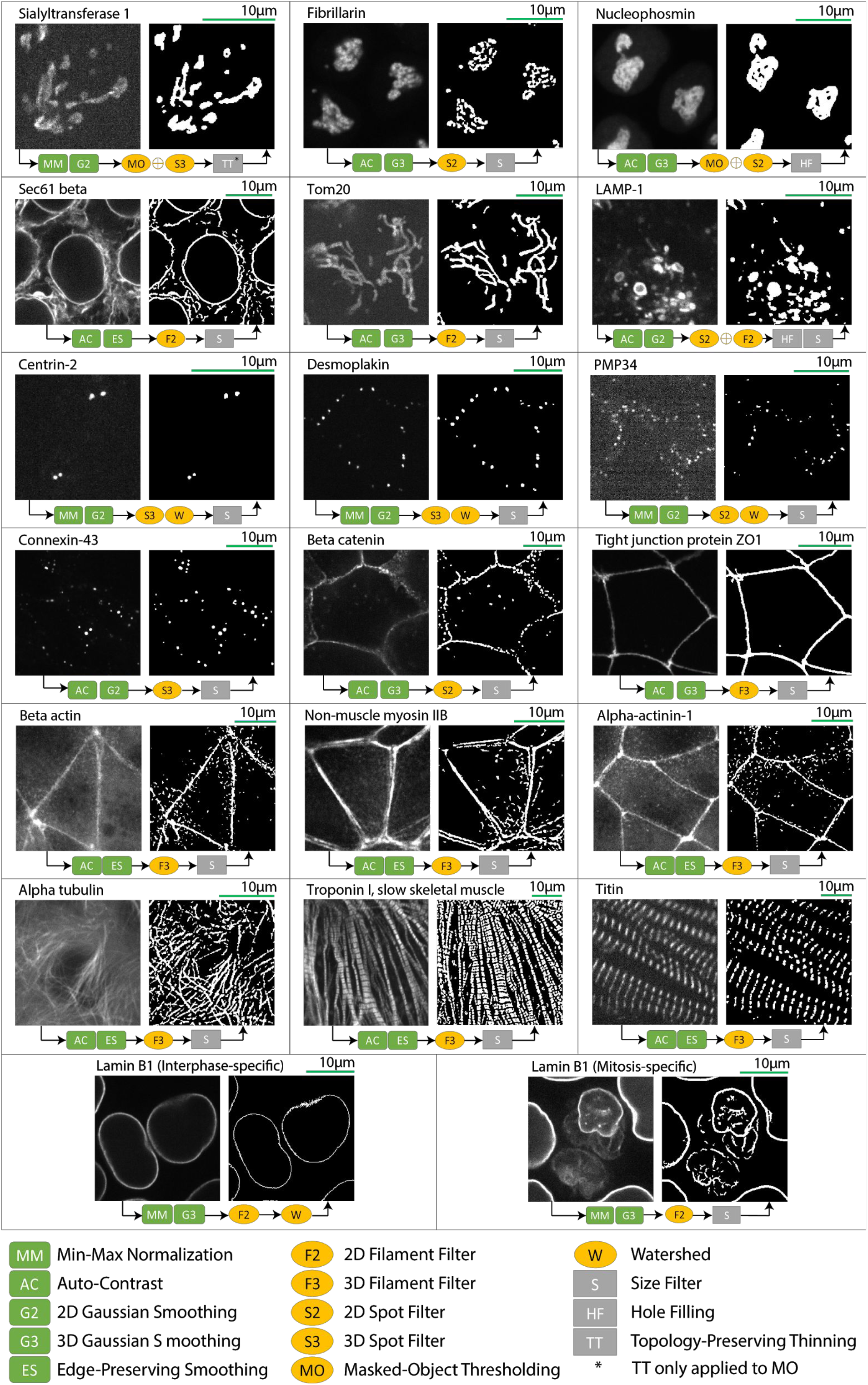
“Lookup table” of classic image segmentation workflows for 20 intracellular structure localization patterns. 18 of the 20 examples consist of image from one tagged protein representing the localization pattern. Examples in the bottom row are both lamin B1 images, but from interphase and mitotic stages of the cell cycle, each one representing a distinct localization pattern requiring a separate segmentation workflow. Each boxed region contains a pair of images with the original image on the left and the result of the classic image segmentation workflow on the right. All original images presented here are single slices from a 3D z-stack of images available online (allencell.org/segmenter). Along the bottom of each image pair is a diagram outlining the steps for that segmentation workflow. The arrows within each diagram represent the transitions between the three steps (pre-processing, core segmentation, post-processing) of the classic image segmentation workflow (**Fig. 1**). Within each workflow step, two symbols directly adjacent to each other represent that the algorithms were applied sequentially while the symbol represents combining the results from both algorithms. The asterisk within the TT symbol for the sialyltransferase 1 workflow indicates that the topology-preserving thinning was only applied to the results from the masked object thresholding algorithm. The target result for LAMP-1 includes filling the larger lysosomes as the protein LAMP-1 labels the lysosomal membrane, but the target structure to detect is the entire lysosome. The specific lookup table in this figure represents our first 20 workflows. As new algorithms are developed for more intracellular structures, the set of algorithms and workflows in the lookup table will be updated on allencell.org/segmenter.

The classic image segmentation and iterative deep learning workflows complement each other – the classic image segmentation workflow can generate sufficiently accurate segmentations for a wide range of intracellular structures for analysis purposes. However, when the accuracy or robustness of the optimal classic image segmentation workflow is insufficient, the iterative deep learning workflow can boost segmentation performance. Conversely, the classic segmentation workflow facilitates the application of deep learning models to 3D segmentation by generating candidate segmentations for an initial ground truth for model training. We have thus developed a new toolkit for 3D fluorescence microscopy image segmentation that (1) is applicable to a wide range of structures, (2) achieves state-of-the-art accuracy and robustness, and (3) is easy to use for cell biology researchers.

### The classic image segmentation workflow

The challenge of designing classic image segmentation algorithms for a large number of distinct intracellular structures led us to a simple 3-step workflow. The steps include a restricted set of image processing algorithm choices and tunable parameters to effectively segment a wide range of structure localization patterns. The classic image segmentation workflow begins with a two-part pre-processing step, intensity normalization and smoothing, followed by the core segmentation algorithms, and ends with a post-processing step. Pre-processing prepares the original 3D microscope images for the core segmentation algorithms. Intensity normalization helps the segmentation be more robust to different imaging inconsistencies, including microscopy artifacts, debris from dead cells, etc., such that the same structures in different sets of images have similar intensity values above background when fed into the core segmentation algorithms (**Fig. 3A**). Smoothing reduces background noise from the microscope and other sources to improve segmentation algorithm performance. The choice of smoothing algorithm depends on the morphology of the intracellular structure (**Fig. 3B**). The core of the classic image segmentation workflow is a collection of algorithms for segmenting objects with different morphological characteristics (**Fig. 4**). This step takes in the pre-processed 3D image stack and generates a preliminary binary segmentation as input into the post-processing step. The final, post-processing step then fine-tunes the preliminary binary segmentations such as by filling holes or filtering by object size, turning them into a final result (**Fig. 5**). The classic image segmentation workflow for a specific structure localization pattern may consist of just one of the core algorithms or it may require a combination of several core algorithms (**Fig. 2**).

**Figure 3:**
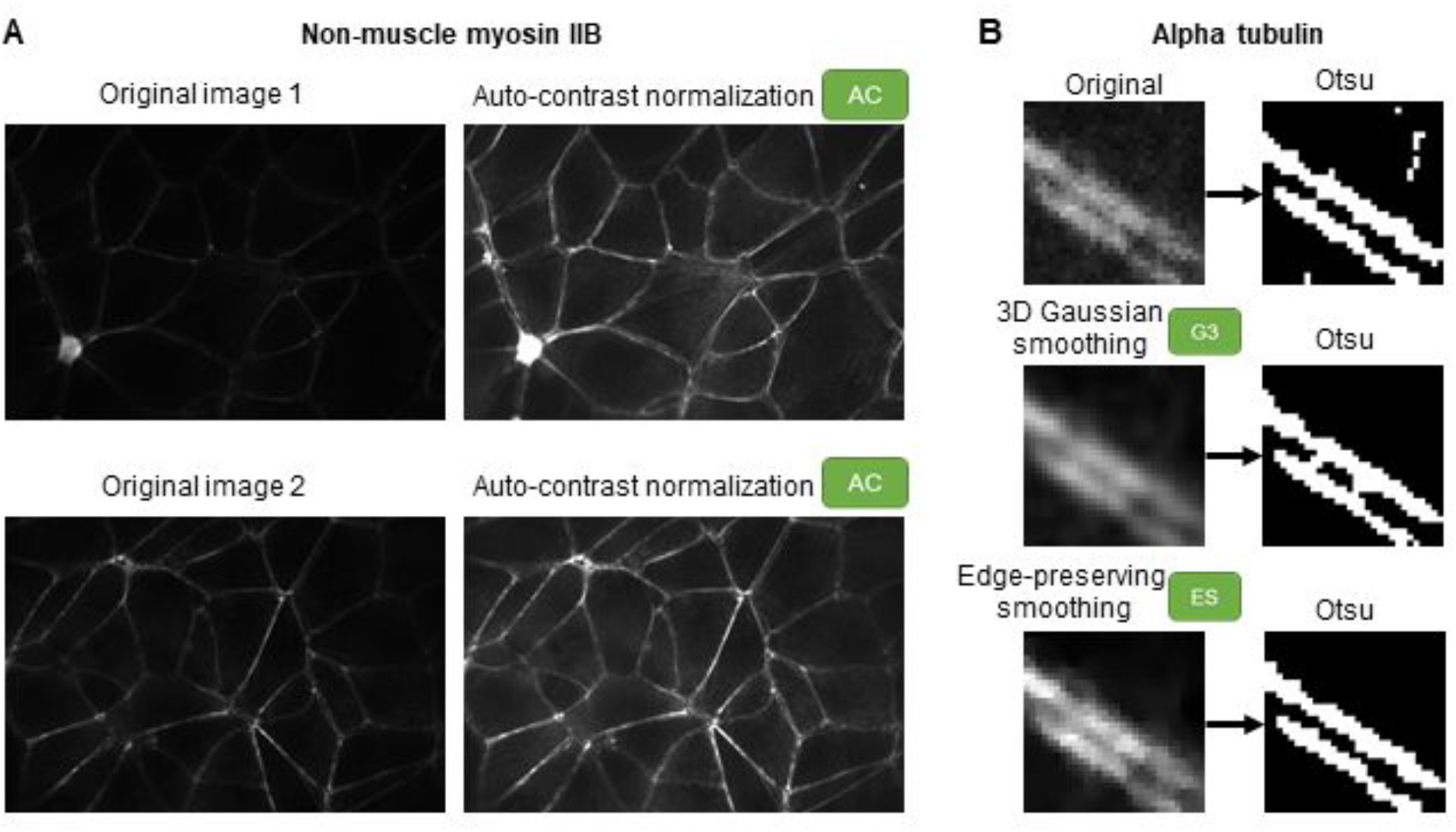
Examples of available pre-processing algorithms in the Allen Cell and Structure Segmenter. **(A)** Two different examples of non-muscle myosin IIB images with corresponding auto-contrast normalization (AC) results. **(B)** An example image of alpha tubulin demonstrating the difference between 3D Gaussian smoothing (G3) and edge-preserving smoothing (ES). The Otsu thresholding results of the original image and the images after two smoothing algorithms are shown to highlight the difference in detecting two parallel tubule bundles.

**Figure 4:**
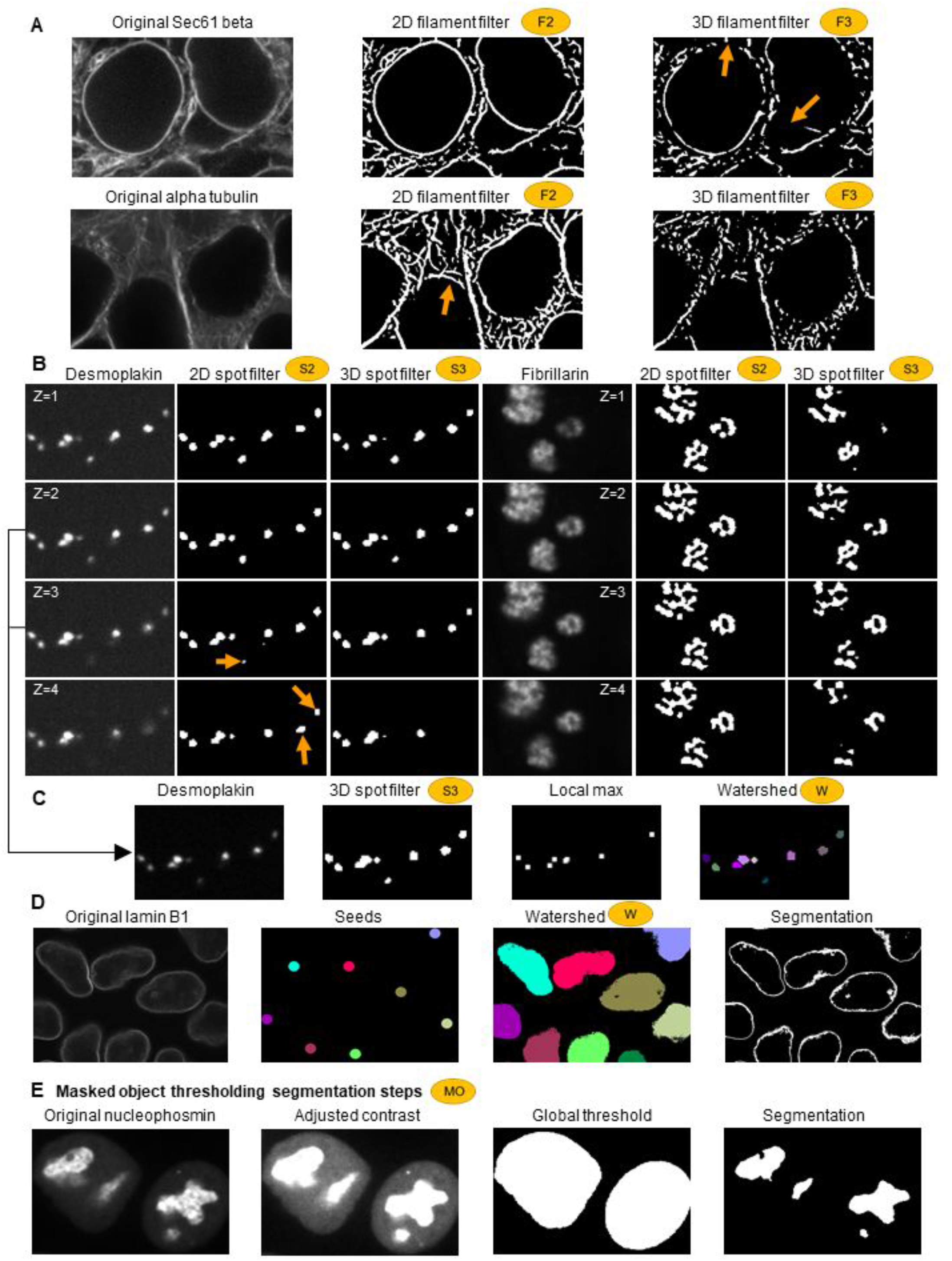
Examples of the available core segmentation algorithms in the Allen Cell and Structure Segmenter. **(A)** Comparison between 2D and 3D filament filters (F2 and F3) applied to Sec61 beta and alpha tubulin images. The binary images are the results of applying a cutoff value (manually optimized for demonstration purpose) on the filter outputs. F2 performs better for curvi-linear shapes in each 2D slice, as seen in Sec61 beta images. F3 performs better for filamentous shapes in 3D, as seen in alpha tubulin images. Orange arrows indicate major segmentation errors. **(B)** Comparison between 2D and 3D spot filters applied to desmoplakin (left three panels) and fibrillarin (right three panels) images. Four consecutive z-slices are shown as four rows. The binary images are the results of applying a cutoff value (manually optimized for demonstration purpose) on the filter outputs. S3 performs better for round, fluorescence-filled shapes in 3D, as seen for desmoplakin images. In this case the S2 filter falsely detects desmoplakin in z-frames where the structure is already out of focus (orange arrows). S2 performs better for localization patterns with a general spotted appearance within each 2D frame, as seen in fibrillarin images. **(C)** Application of the watershed algorithm (W) on S3 segmentation of desmoplakin to separate merged spots. Left to right: The maximum intensity z-projection of the middle two z-slices of desmoplakin in **(B)** and the corresponding S3 result, the detected local maximum (small white dots) and the final results after applying the watershed (W) algorithm with the local maximum as seeds. **(D)** Application of the watershed algorithm (W) directly on pre-processed lamin B1 interphase images for segmentation with automatically detected seeds. The watershed line of the watershed output is taken as the segmentation result. **(E)** Application of the steps of masked object thresholding (MO) to nucleophosmin images. From left to right: an original image of nucleophosmin, the same image with adjusted contrast to highlight lower intensity signal throughout the nucleus, the result of a global threshold obtaining the overall shape of nuclei used as a mask, and the result of the Otsu threshold within individual nuclei as the final segmentation image. Compared to traditional global thresholding, masked-object thresholding is more robust to variations in intensity between different nuclei in the same image.

**Figure 5:**
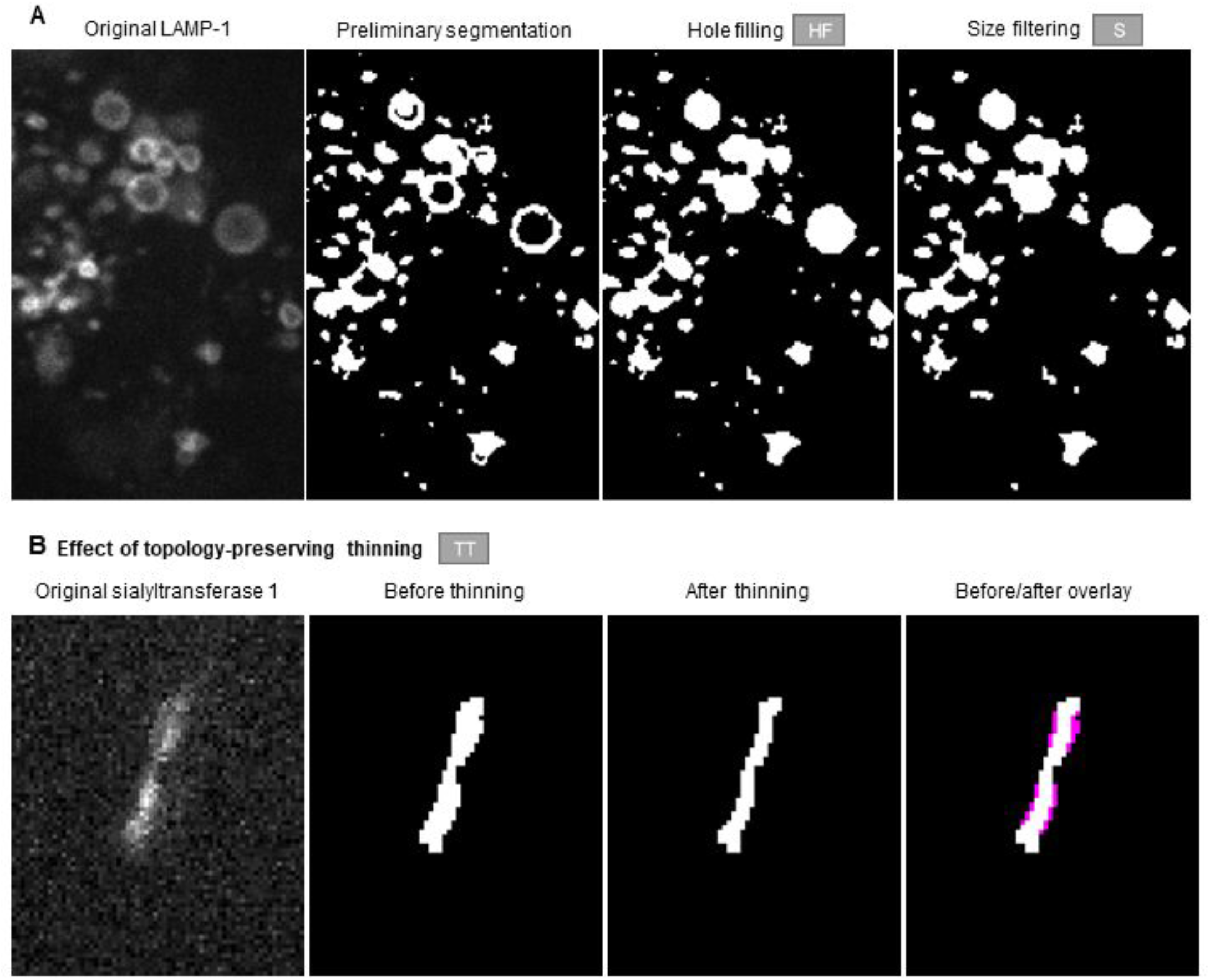
Examples of the available post-processing algorithms in the Allen Cell and Structure Segmenter. (**A**) Demonstration of the hole-filling algorithm (HF) and the size filter (S) applied to the preliminary segmentation results of LAMP-1 images. The target result for LAMP-1 includes filling the larger lysosomes as the protein LAMP-1 labels the lysosomal membrane, but the target structure to detect is the entire lysosome. (**B**) Demonstration of the topology-preserving thinning algorithm (TT) on sialyltransferase 1 images. The magenta in the last column represents pixels that are removed after applying the TT algorithm.

### Application of the classic image segmentation workflow to segmentation of over 30 intracellular structure localization patterns

We applied the classic image segmentation workflow to 3D images of over 30 fluorescently tagged proteins, each representing different intracellular structures. Structures were imaged in two different cell types, the undifferentiated hiPS cell and the hiPSC-derived cardiomyocyte. The tagged proteins representing these structures exhibited different expression levels and localization patterns in these two cell types. Certain structures also varied in their localization patterns in a cell cycle-dependent manner. Together, this led to over 30 distinct intracellular structure localization patterns, which we used to develop and test the classic image segmentation workflow. A key decision point for any segmentation task is the targeted level of accuracy. This is a function of several factors including: the size of the structure, the limits of resolution and detection for that structure, the goal of the subsequent analysis, and the effort required to obtain any given target accuracy. In general, we aimed to be consistent with observations in the literature about the structure and to obtain segmentations useful for 3D visualization.

For example, our alpha tubulin segmentation workflow (**Fig. 2**) describes where the microtubules primarily localize and is detailed enough to generate a reasonable 3D visualization, but does not take known structural properties, such as the persistence length of microtubules, into account [Gan et al., 2016]. We also addressed the blurring of the boundaries of the structure arising from the resolution limits of fluorescence microscopy. Depending on both the contrast setting of the image and the parameters of a given segmentation algorithm, the resultant binary image can vary significantly. For example, segmentation of a mitochondrial tubule can result in segmented tubules of varying width (**Fig. 6**). To establish a consistent baseline of how to detect the blurred boundary, we used a fluorescently tagged mitochondrial matrix marker as a test structure and picked the segmentation parameter that most closely matches EM-based measurements of mitochondria in human stem cells ([Bukowiecki et al. 2014, Niclis et al. 2015]; see methods). We then used the resultant combination of contrast settings and object boundary setting as a consistent target for the creation of the other intracellular structure segmentation workflows.

**Figure 6:**
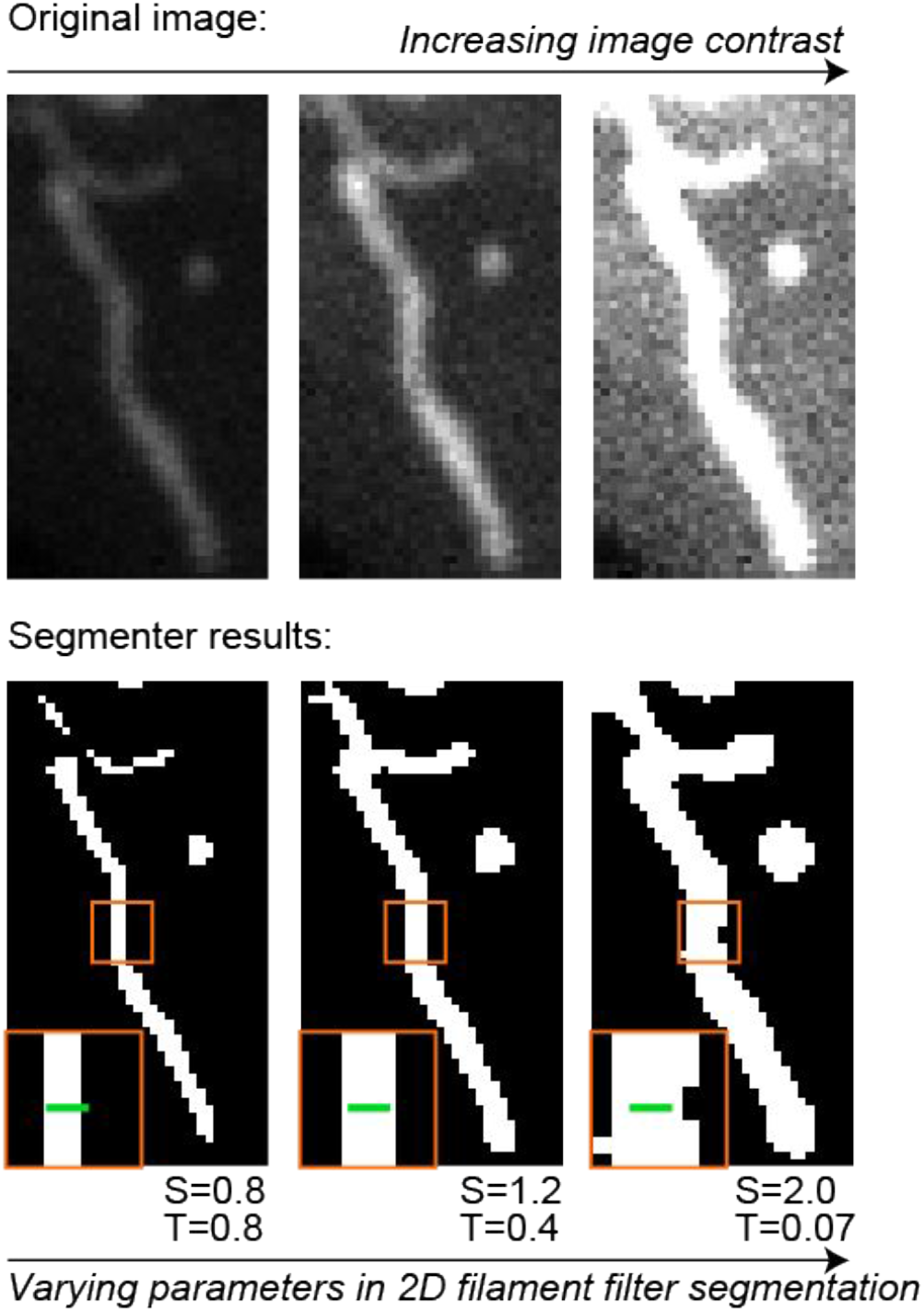
Comparisons of detected object boundaries dependent on segmentation parameters. To identify a consistent target for detecting boundaries of the many different tagged intracellular structures, we demonstrate the effect of different segmentation parameters on the segmentation of mitochondrial tubules marked with a matrix-targeted mCherry in transiently transfected hiPS cells. The top row shows the same tubule with increasing brightness and contrast settings from left to right for visualization purposes only. The bottom row shows the result of increasing the kernel size parameter (S) while decreasing the threshold parameter (T) in the 2D filament filter segmentation algorithm, which was applied to this image. The brightness and contrast settings in the top row are set to match the segmentation results in the bottom row to demonstrate that each of the segmentation results are reasonable given the input image. The green line within the inset in the bottom row is 260 nm, the diameter of mitochondria based on EM images of human stem cells. The second column represents the segmentation result that the collection of classic image segmentation workflows in the look-up table aimed to consistently target for each intracellular structure localization pattern.

Application of the classic segmentation workflow with these structure target criteria culminated in the intracellular structure look-up table of segmentation workflows for 20 selected distinct structure localization patterns (**Fig. 2)**. These classic image segmentation workflows significantly improved segmentations compared with a set of 22 common baseline algorithms, including both global and local thresholding (**Fig. 7**). We found that for structures with similar morphological properties, the same series of algorithm choices resulted in successful segmentation (compare workflow diagrams in rows 5 and 6 in **Fig. 2**). However, the parameter values for the best result still varied. The structure segmentation look-up table and accompanying 3D z-stack movies (**Fig. 2** and allencell.org/segmenter) thus serve as a guide for which segmentation workflow (set of algorithm steps and parameter values) is a good starting point for a user’s particular segmentation task. The classic image segmentation workflow component of Segmenter is openly accessible at allencell.org/segmenter. For each example in the lookup table, the specific workflow and accompanying algorithm parameters, fine-tuned on our data, are preset in a workflow-specific Jupyter notebook and accompanying “pseudocode” (see Supplemental Information) for rapid referencing and initial testing. If the preset parameters do not generate satisfactory segmentation results on user data, the parameters can be adjusted and assessed directly within the Jupyter notebook via an embedded 3D viewer. Step by step suggestions of which parameters to adjust are also included to help the user.

**Figure 7:**
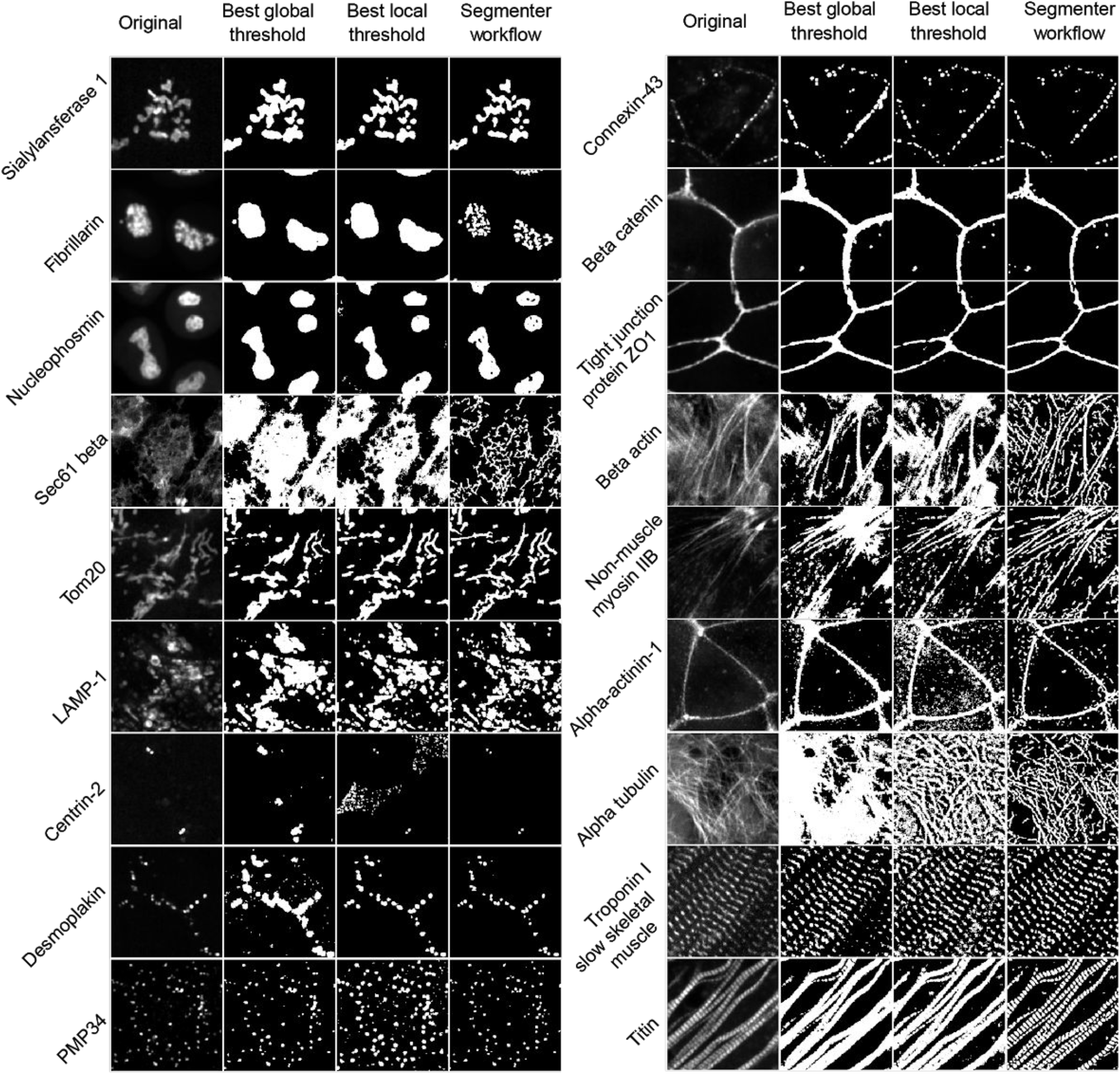
Comparison of segmentation results between the Allen Cell and Structure Segmenter (Segmenter) classic image segmentation workflow and 14 global and 8 local thresholding algorithms applied to the 20 structure localization patterns in the lookup table (**Fig. 2**). All images correspond to the maximum intensity projection of a z-slice of choice plus and minus one z-slice. The first column represents the original image. The second, third and fourth columns display the optimal global algorithm result, optimal local algorithm result, and the Segmenter classic image segmentation workflow result, respectively (**Fig. 2;** see methods for list of algorithms tested).

### The iterative deep learning workflow

The aim of the iterative deep learning workflow (**Fig. 1B**) is to improve segmentation accuracy and robustness for situations where the classic image segmentation workflow is insufficient. It applies the concept of incremental learning [Schlimmer et al., 1986] to iteratively improve segmentation results. The segmentation results from the classic image segmentation workflow provide us with a set of 3D segmentation images that have the potential to be used as a ground truth for deep learning purposes. However, the accuracy of these results is often not uniform for all images. Instead, the greatest accuracy is generated in subsets of images or specific regions within images. We developed and tested two human-in-the-loop strategies, sorting and merging, to convert a set of classic image segmentation workflow results into an acceptable 3D ground truth image set for model training. These straightforward human-in-the-loop strategies do not involve any manual painting of the 3D structure, but still incorporate human knowledge into curating high-quality segmentation ground truth images. These first curated results from the classic image segmentation workflow can then be used as a starting point to train the first model. The segmentation results of the first model can once again be curated to provide a second, improved ground truth data set to create a second, improved segmentation model and so on for any number of iterations. Iterations can also be performed by combining segmentation results from several deep learning models via the merging and sorting strategies to further improve the resultant model. This approach thus eliminates the need for a large set of training data. The iterative deep learning workflow is implemented in an easily accessible way and with minimal tunable parameters. Specifically, users put raw images and training ground truths (segmentation images) in the same folder following a prescribed naming convention and set a few parameters that vary depending on image resolution and imaging modality. The details of building models, setting hyper-parameters, training the models, and so on, are handled automatically in a way that is designed and optimized for 3D fluorescence microscopy images.

### Application of the iterative deep learning workflow generates an accurate and robust lamin B1 segmentation

Image-to-image variation and cell-to-cell variation are two common scenarios in which the classic image segmentation workflow may not be sufficiently robust to data variation. For example, algorithms in the classic segmentation workflow do not always handle the image-to-image variation that arises within a dataset due to differences in biological and/or microscopy imaging conditions. In this case, a simple approach to preparing a ground truth segmentation image set is to sort segmented 3D images into “accept” or “reject” categories and only use the accepted images for initial model training (**Fig. 8**). The subsequent deep learning model may end up more robust to image-to-image variation because it incorporates contextual knowledge, which the classic segmentation workflow algorithms are incapable of doing. Similarly, within the same image, cells at different stages of the cell cycle may exhibit distinct structure morphologies or fluorescence intensities since interphase and mitotic structures can differ dramatically. In this case, two (or more) different segmentation parameters or algorithms might permit both types of structure localization patterns to be well-segmented, but two sets of parameters or algorithms cannot normally be applied to the same image. With the aid of a simple image editing tool, such as those available through ImageJ or ITK-SNAP, however, specific regions of interest within an image can be manually circled and masked. Different parameter sets or algorithms are then applied to each of these regions, and the results merged into one single segmentation ground truth for that image (**Fig. 9**). A single deep learning model usually has sufficient representation capacity to learn all such variations within the image.

**Figure 8:**
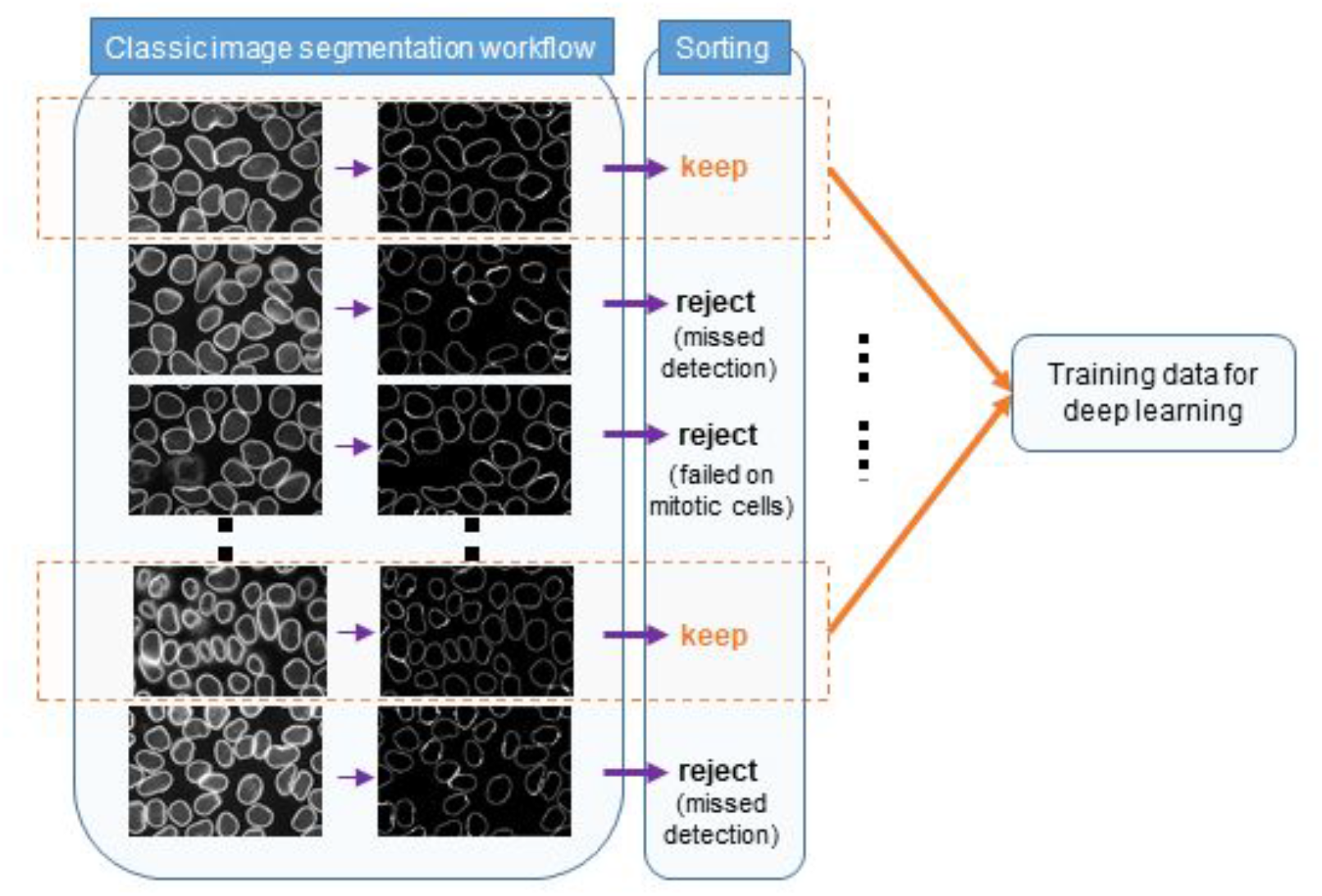
Schematic of the human-in-the-loop sorting strategy for generating ground truth data sets from classic image segmentation workflow results for training subsequent 3D deep learning models. A classic image segmentation workflow was applied on a set of lamin B1 images, and then the segmented images were manually curated to sort out the images to keep. These images directly become the 3D deep learning training set.

**Figure 9:**
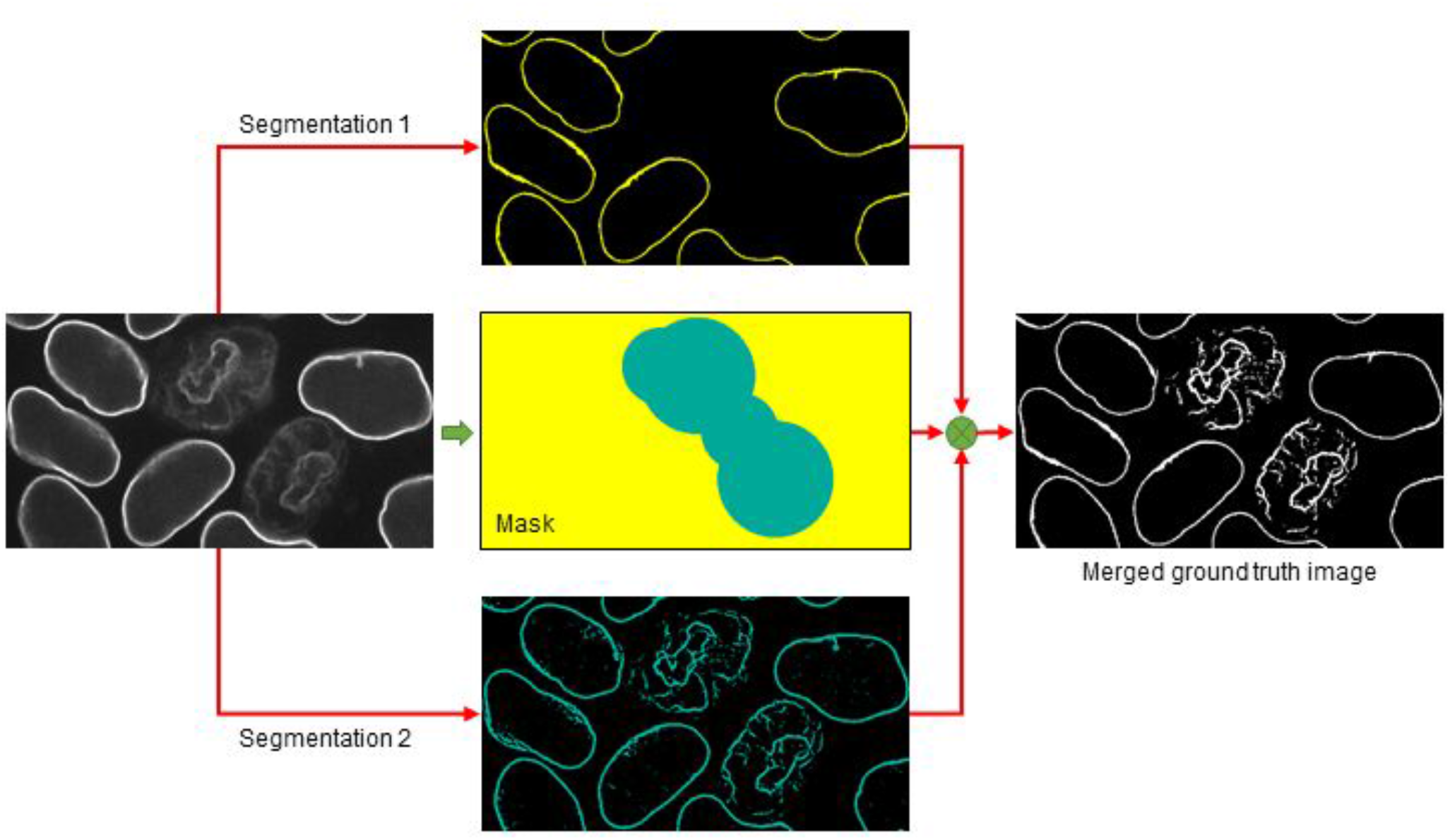
Schematic of the human-in-the-loop merging strategy for generating ground truth data sets from classic image segmentation workflow results for training subsequent 3D deep learning models. Two different classic image segmentation workflows were applied to the same lamin B1 images. One workflow worked well on interphase lamin B1 localization patterns (yellow) and the other worked better on mitotic lamin B1 localization patterns (cyan). A mask made up of four circles with two different radii was manually created in Fiji with the “PaintBrush” tool. A single ground truth image was then merged (⊗) based on the two segmentations by taking the segmentation displayed in yellow within the yellow area and the segmentation displayed in cyan within the cyan area on the mask. This merged ground truth image is used as part of the 3D deep learning training set.

To demonstrate the segmentation accuracy and robustness achievable by the iterative deep learning workflow, we applied this workflow to the segmentation of lamin B1 images (**Fig. 10**). The lamin B1 localization pattern changes dramatically through the cell cycle, changing from a thin shell around the nucleus in interphase to a variable, wavy pattern during mitosis. This significant difference in localization patterns created a challenge for a classic image segmentation approach. To address this, we first built a classic image segmentation workflow to segment lamin B1 in interphase cells (**Fig. 2**). The best core algorithm to obtain accurate segmentation of this lamin B1 shell depended on generating an automatic seed in the center of each nucleus for the subsequent watershed algorithm. This automatic seeding was performed on the center slice of the 3D image stack, where most of the nuclei were easily detectable. However, this automatic seeding step sometimes failed, especially for cells with nuclei positioned above or below this center slice (blue arrows in **Fig. 10**). Further, as expected, mitotic cells in the image were not successfully segmented with this lamin B1 interphase-specific segmentation workflow (yellow arrows **Fig. 10**). From an initial set of 80 segmented images, only eight fell into the “keep” sorting category. The rest of the images either failed in the automatic seeding step or contained mitotic cells, which could therefore not be used as a ground truth. However, a first iteration deep learning model based only on these eight image stacks as its initial ground truth generated lamin B1 segmentations that picked up all interphase nuclei in all 80 images. This was an improvement from 85% to 100% when considering individual interphase nuclei and an improvement from 20% to 100% when considering entire image fields (e.g. an image field can fail if a single nucleus is not detected). In the second iteration of this workflow, we manually circled mitotic cells in 22 additional images and applied a separate classic image segmentation workflow for mitotis-specific lamin B1 localization patterns (**Fig. 2**). We then merged the interphase lamin B1 segmentation results from the first deep learning model with the mitotic lamin B1 segmentations obtained from the classic workflow to build a new training set out of these 22 images. This second iteration of a lamin B1 segmentation deep learning model generated successful lamin B1 segmentations for all interphase and mitotic cells in the original 80 images.

**Figure 10:**
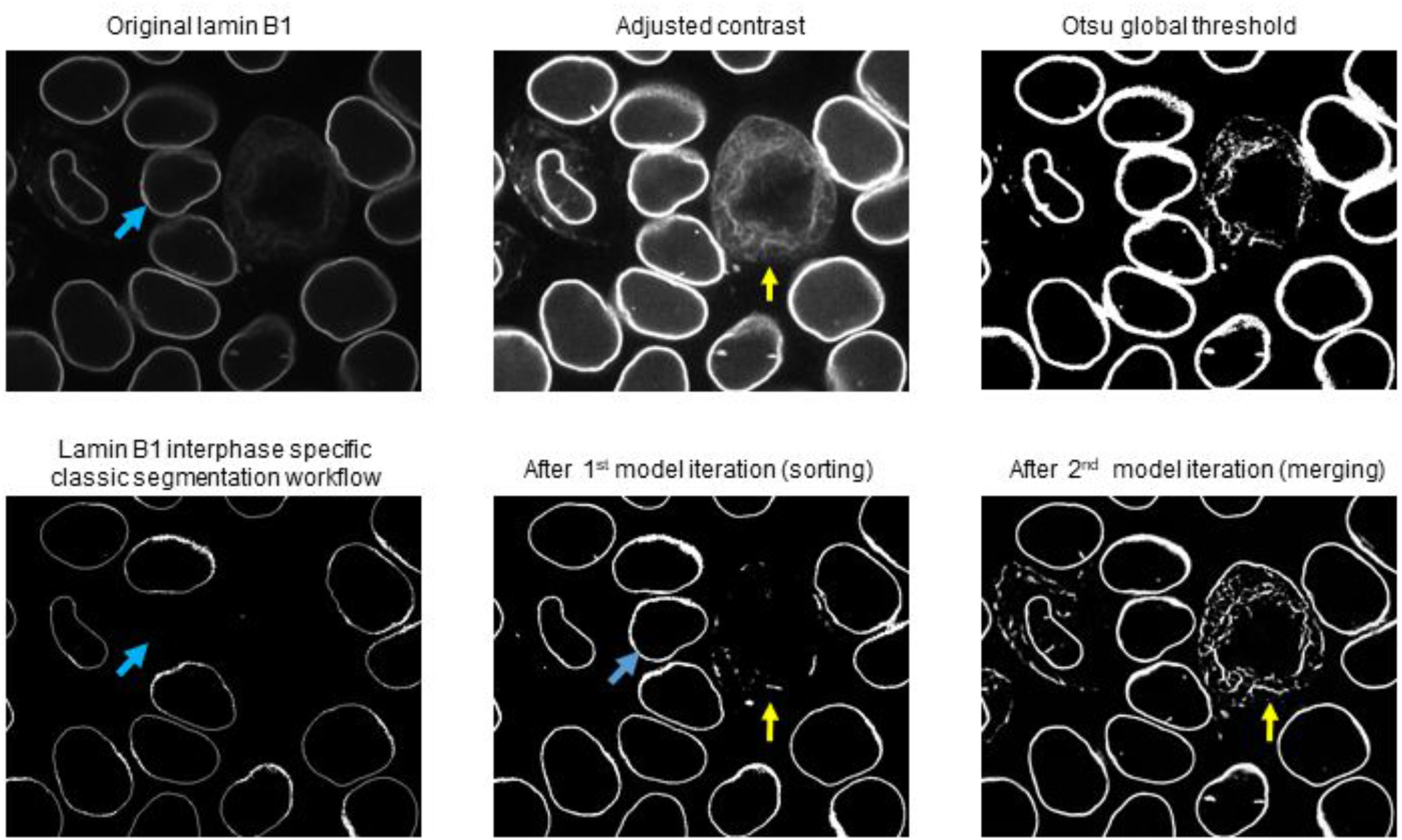
Application of the iterative deep learning workflow to generate robust lamin B1 segmentations. The first two images in the top row represent a middle z-slice from the original lamin B1 image and a second version of the same image with adjusted contrast settings to highlight the fine structure in the lamin B1 mitotic localization pattern (yellow arrow). The third column in the top row represents the result of a standard Otsu thresholding segmentation as a baseline. The first image in the bottom row shows the Segmenter classic image segmentation workflow result. The middle image shows the result after the first iteration of the deep learning model trained by a ground truth set generated by sorting. Finally, the third image shows the result after the second iteration of the deep learning model trained on a ground truth set generated by merging. The blue arrow indicates an interphase lamin B1 localization pattern that was originally missing in the classic image segmentation workflow but was detected using the iterative deep learning models. The yellow arrow indicates a mitotic lamin B1 localization pattern that was detected after the second iteration of the deep learning model.

It is worth mentioning that applying a model trained from an iterative deep learning workflow on a new image will output an image where each pixel has a value between 0 and 1 indicating the probability of being part or not part of the target structure. A cutoff value needs to be applied to generate the binary segmentation. We used a default cutoff of 0.5, which works fine in most cases. The cutoff value can also be selected automatically when a segmentation target is available. In this case a cutoff value can be identified that achieves the highest score on any particular desired segmentation metric (e.g., mean intersection over union). The Segmenter is a powerful toolkit for the 3D segmentation of intracellular structures in fluorescence microscope images. The Segmenter combines a streamlined collection of selected standard and cutting-edge classic image segmentation algorithms with a suite of preset classic image segmentation workflows for 20 distinct intracellular structure patterns and with a novel iterative deep learning segmentation workflow. The classic image segmentation workflow together with the lookup table should provide users with a straightforward starting point for their own basic segmentation needs. More challenging segmentation tasks can benefit from the complementary approach of the classic and the iterative deep learning segmentation workflows. This combined approach permits training of deep learning models that can successfully segment different structure localization patterns within a single image. Two of the most significant challenges to creating robust useful segmentation methods for 3D quantitative analysis of cell behavior is to detect all instances of the cell or structure within the entire image and to do so successfully over time, especially if the cell or structure changes [Ulman et al., 2017]. This is required, for example, to capture the dynamic behavior of neighboring cells or neighboring structures within cells. The challenge here is that any cell or structure missed within one image or within one timepoint of a timeseries significantly reduces the analyzable data set. The success of our joint classic and iterative deep learning approach in improving the detection of all nuclei in a lamin B1 image from 20 to 100% with just one iteration of the model suggests great potential of applying this approach to the robust, automated segmentation of entire images of cells and intracellular structures in time.

### Training Assays for semantic segmentation of nuclei/mitotic DNA

In 3D fluorescent spinning disk confocal microscopy, a cellular structure imaged as a solid shape, like the DNA representing the filled nucleus when cells are labeled with DNA dye, will have blurry boundaries due to the diffraction of light, which is especially limiting in the axial z direction. As a consequence, locating an accurate and consistent boundary of a filled fluorescent nucleus merely from the fluorescent images is challenging. For this reason, we propose to leverage specialized biological experiments to obtain a preliminary nuclear segmentation with a more biologically correct boundary. Starting from this preliminary segmentation, we can then use iterative deep learning to build a deep neural network model that can segment nuclei from 3D microscopy images with more accurate boundaries than are visible in the assay data the model is applied to.

Our target problem is the 3D semantic segmentation of the nuclei of interphase cells and the mitotic DNA during mitosis (representing the “nucleus” during nuclear envelope break-down) from DNA dye images. We use “nuclei” to refer to both nuclei in interphase cells and reforming nuclei in the two daughter cells in telophase/cytokinesis. We use “mitotic DNA” to refer to the “nuclei” in prophase, metaphase and anaphase. Semantic segmentation can be defined as the overall assignment of each voxel in an image stack as being or not being part of one type of foreground objects (nuclei/mitotic DNA in this case), without identifying instances (e.g., individual nuclei). We used images from two separate secondary assays for this Training Assay approach: 1) mEGFP-tagged lamin B1 (labels the nuclear envelope) for interphase nuclei and 2) mEGFP-tagged H2B (labels a subset of DNA histones) from the hiPSC ingle-Cell Image Dataset. In each image, a field of view (FOV) of cells is imaged in four different channels: DNA dye, mEGFP-tagged lamin B1 (or mEGFP-tagged H2B), cell membrane dye, and bright-field transmitted light. Lamin B1 labels a subset of the nuclear lamina and is located just inside the inner nuclear envelope. For hiPS cells during interphase, the segmentation target is the equivalent of a filled lamin B1 shell representing the nucleus (**Fig. 11**). During mitosis, the lamin B1 signal re-localizes to other regions of the cell as the nuclear envelope partially breaks down. Therefore, during mitosis, the segmentation target is the condensed mitotic DNA, which then decondenses at the end of mitosis as the nuclei reform. Mitotic DNA is visible via the DNA dye but has much more consistent and higher image quality when imaged via endogenously tagged mEGFP-tagged H2B (bottom row of **Fig. 12**), which we thus use H2B as a “cleaner” equivalent for visualizing mitotic DNA during mitosis.

**Figure 11:**
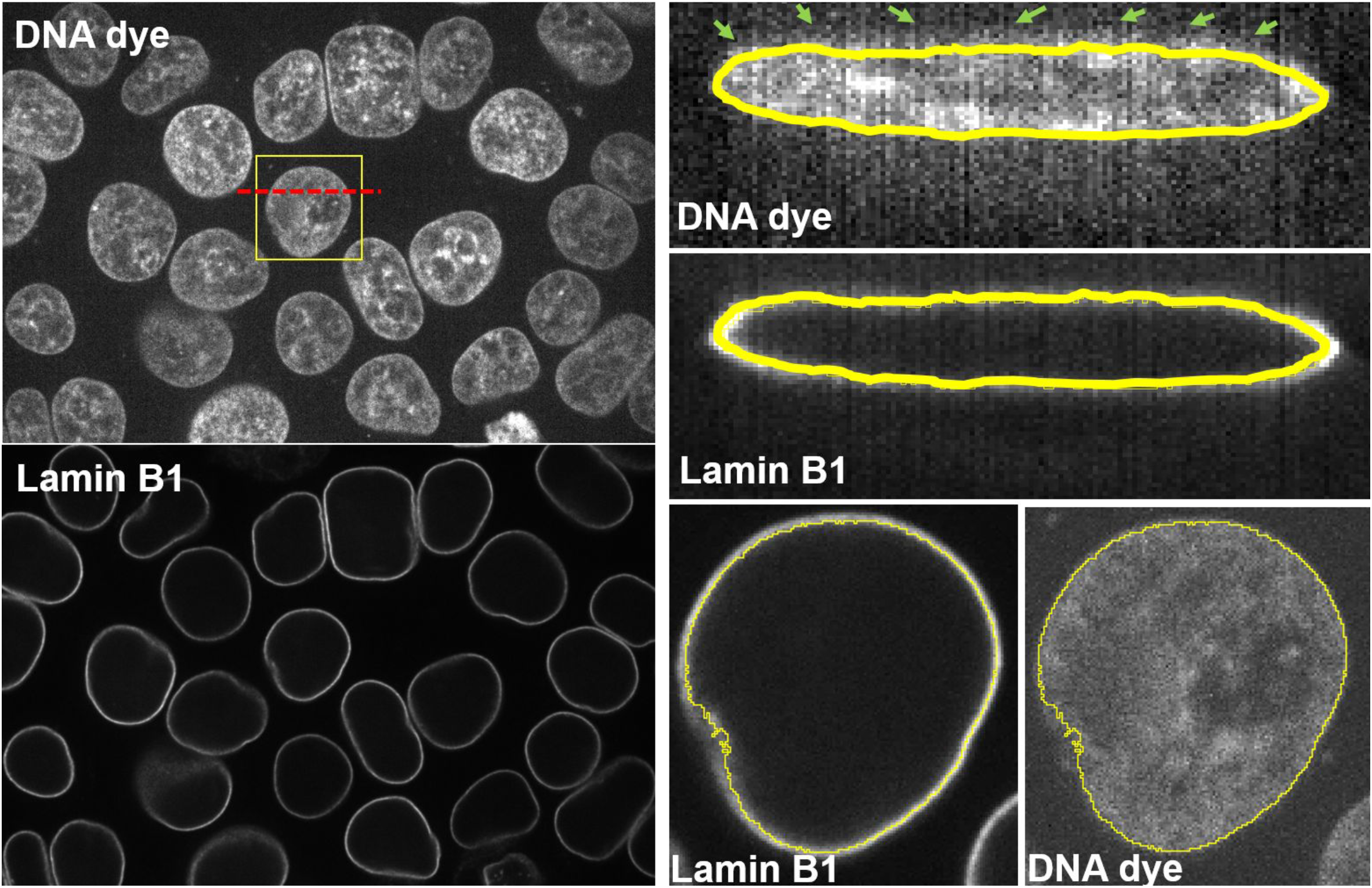
Illustration of the lamin B1 training assay. Left: An example z slice of a multi-channel 3D microscopy showing both DNA dye (top left panel) and mEGFP-tagged lamin B1 (bottom left panel) of the same field-of-view. Right: Zoomed-in images of the nucleus in the yellow box in the DNA dye panel. Top two panels show orthogonal views lamin B1 and DNA dye images along the red dash line and bottom two panels show the front view. The lamin B1 segmentation result from the classic image segmentation workflow is shown as the yellow line overlaid onto both the lamin B1 image and DNA dye image, in order to demonstrate the correspondence between lamin B1 and the nuclear boundary. Due to optical properties of the microscope, including diffraction of light, especially along the z direction, it is not straightforward to identify the exact boundary of the nucleus from the DNA dye (see the green arrows). However, the nuclear envelope can be visualized via the tagged lamin B1, which creates a “shell” around the nucleus. This thus permits a more biologically accurate detection of the nuclear boundary, especially in the z direction. This training assay-derived boundary can then be used to train a deep learning model to segment interphase nuclei from DNA dye with the added accuracy from the lamin B1 assay.

**Figure 12:**
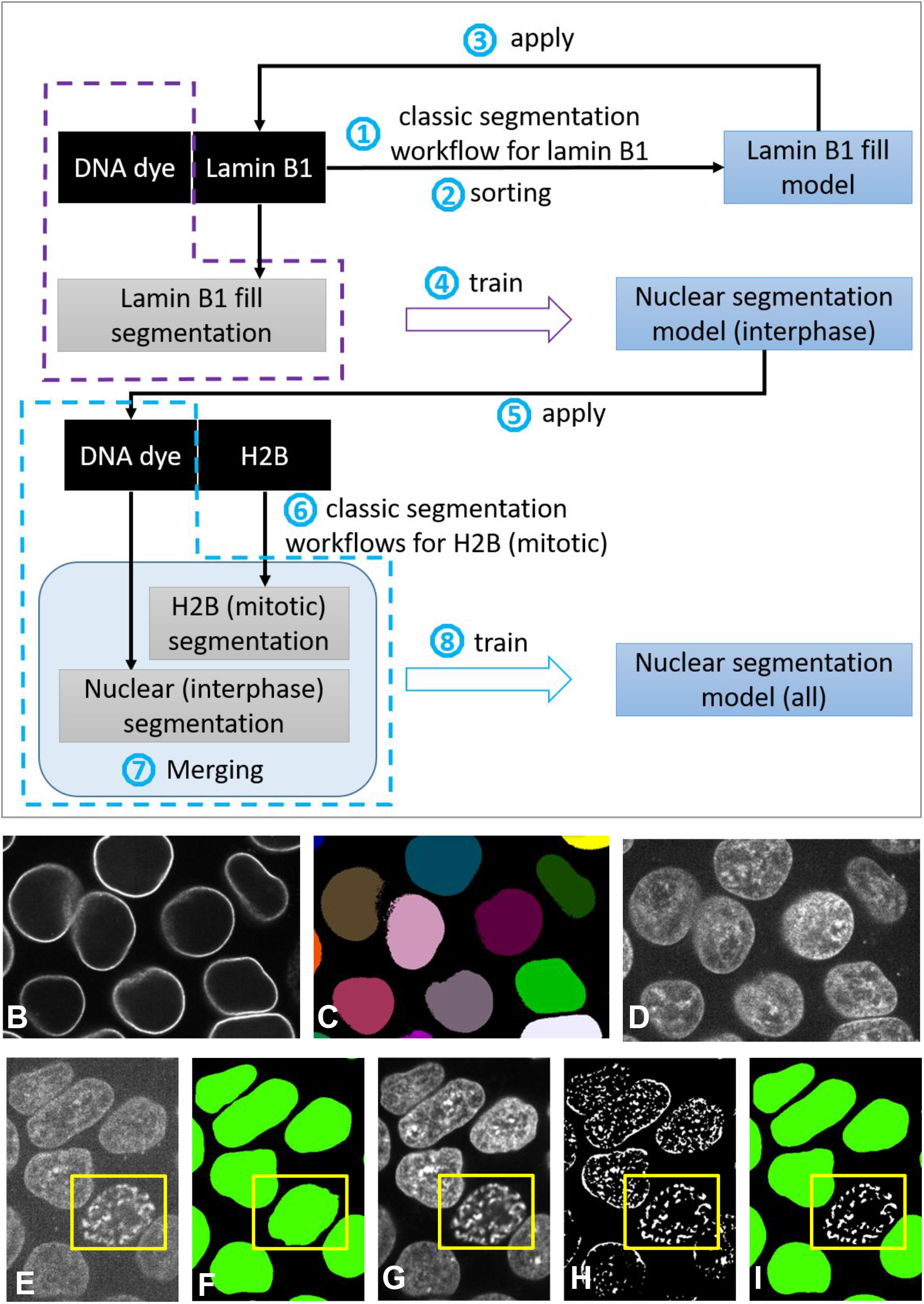
Overview of the computational steps for training a DNA dye-based nuclear segmentation model for segmenting nuclei/mitotic DNA from 3D images of DNA-dye labeled hiPSCs. Panel A outlines the eight main steps, as marked by the numbers in blue circles. Step (1) and (2) create the lamin B1 fill model using the same iterative deep learning workflow and data as illustrated in **Fig. 8**, except using the filled shell of lamin B1 segmentation as the segmentation target for training. In step (3), the lamin B1 fill model is applied to the lamin B1 channel (see B) of the multi-channel image stacks from the lamin B1 cell line. The generated lamin B1 fill segmentation (see C) is used in step (4) to train the interphase nuclear segmentation model that takes DNA dye channel (see D) of the same multi-channel image stacks as input. In step (5), the DNA dye-based interphase nuclear segmentation model is applied the DNA dye channel (see E) of the multi-channel image stacks from the H2B cell line. The result (see F) contains good segmentation for interphase nuclei, but not ideal for the one in the yellow box, which already starts the mitosis. In step (6), we apply classic segmentation workflows for mitotic H2B (one workflow for prometaphase/metaphase and one workflow for other mitosis stages) on the H2B channel (see G) of the same image stacks to obtain mitotic specific H2B segmentation (see H). In step (7), the interphase nuclear segmentation (F) and the mitotic H2B segmentation (H) are merged using the merging curator of Segmenter to create a merged ground truth (see I), which is used in step (8) to train the overall DNA dye-based nuclear segmentation model.

### Training Assay for semantic segmentation of the cell membrane

The signal from the plasma membrane dye used to label the boundaries of all the cells in this hiPSC image dataset suffers from photobleaching even during a single z-stack acquisition via 3D spinning disk confocal microscope. All cells are imaged from bottom to top, and thus the cell membrane at the top of the cells often shows very weak signal, due to both dye labeling of a very thin membrane and the photobleaching. A weak signal at the top of the cells makes accurate segmentation of the cell membrane via only the dye very challenging.

Our target problem is the 3D semantic segmentation of the cell membranes from membrane dye images. The secondary experimental assay for this Training Assay approach consists of using the CAAX cell line from the hiPSC Single-Cell Image Dataset. This cell line contains the membrane-targeting domain of K-Ras tagged with mTagRFP-T. As shown in Fig. 13 (see the blue arrows), the signal to noise ratio (SNR) of the cell membrane top in the CAAX channel is much stronger than in the membrane dye channel. This permits us to generate more accurate segmentations of the cell membrane at the top of cells via the CAAX channel and then impose this segmentation result as the ground truth for training a deep neural network for the segmentation of the cell membrane via the membrane dye.

**Figure 13:**
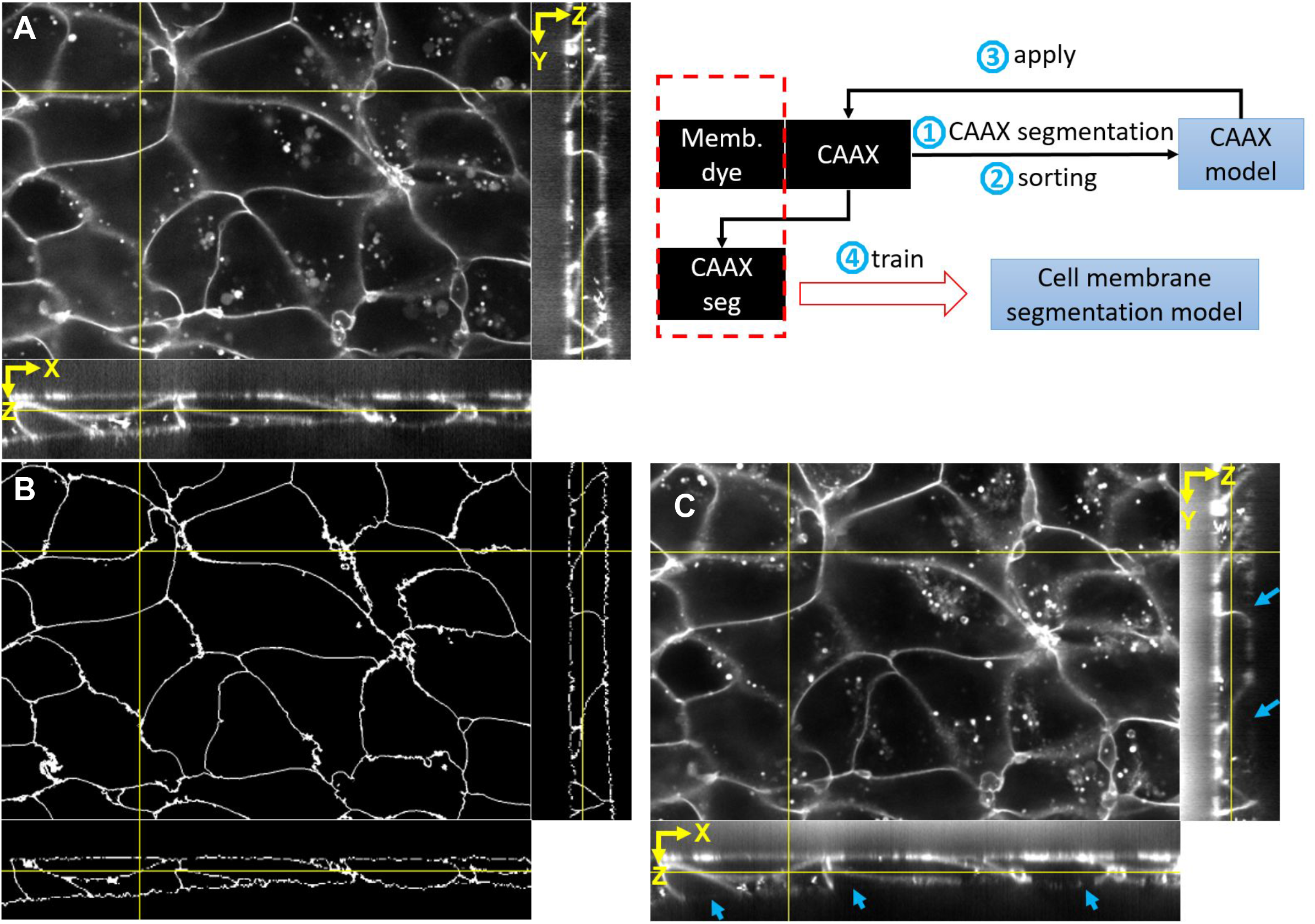
Overview of the computational steps for training a membrane dye-based cell segmentation model for segmenting the boundaries of cells from 3D images of membrane-dye labeled hiPSCs. The upper right panel outlines the four main steps, as marked by numbers in blue circles. In step (1), we apply a watershed-based algorithm on the CAAX (endogenously tagged cell membrane) images to obtain initial CAAX segmentation results. See the method section for details about the CAAX segmentation algorithm. In step (2), we use the sorting curator in Segmenter to go through the initial CAAX segmentation results to find successful ones to train a CAAX segmentation model. In step (3), we apply the CAAX segmentation model on the CAAX channel (see A) of the multi-channel image stacks from the CAAX cell line to obtain CAAX segmentations (see B), which are used in step (4) to train the cell membrane segmentation model that takes cell membrane dye (see C) of the same multi-channel image stacks as input. From the orthogonal views (YZ and XZ) in A-C, we can observe that the CAAX image (A) has much better signal near the top of the cells and therefore can help generate more accurate cell boundary segmentation target (B), comparing the membrane dye (C; as marked by the blue arrows).

### Going from segmentation “models” to segmentation “algorithms”

The ML based segmentation methods we presented so far are all “models”, which take a single-channel image as input and predict the probability of each voxel of being part of the segmentation target or not, which can be then binarized into the segmentation by applying a cutoff (see “***Application of the iterative deep learning workflow generates an accurate and robust lamin B1 segmentation****”*). Beyond basic “models”, we can develop segmentation “algorithms”, which may take a single-channel or multi-channel image as the input, employ one or more “models”, and the model outputs are then processed through one or more steps to generate the final segmentation result. In the sections below, we introduce two such algorithms: (1) a DNA and membrane dye-based 3D cell and nuclear segmentation algorithm, which generates 3D instance segmentations of cells and nuclei from a multi-channel image with both DNA dye and membrane dye; (2) a lamin B1 segmentation algorithm, which takes both the lamin B1 channel and membrane dye channel as input and generates more biologically accurate lamin B1 segmentation than the basic lamin B1 model developed from the iterative deep learning workflow.

### 3D instance segmentation of cell and nucleus

With the three aforementioned Training Assays, we can build two core ML segmentation models: **a membrane dye-based cell segmentation model** and a **DNA dye-based nuclear segmentation model** using the Segmenter (see Methods for details and Fig. 14 for overview). To convert the binary semantic segmentations of the entire FOV into individual cell and nuclear objects we required two additional deep learning models: 1) a seeding model (minor variation from the nuclei segmentation model) and 2) a cell pair detection model (a Faster-RCNN based object detection model, see Methods for details). Combining all these models, we can develop a **DNA and membrane dye-based 3D cell and nuclear segmentation algorithm**. Briefly,we use the membrane dye-based cell segmentation model output to cut the seeding model output and generate one seed for each cell, which we then use in a seeded watershed performed on the membrane dye-based cell segmentation model output to generate instance segmentation of cells. Then, cell instance segmentations are propagated to the DNA dye-based nuclear segmentation model output to generate the instances of nuclei/mitotic DNA. Because the two daughter cells after mitosis (before completely entering interphase) should still be considered as one single “biological object” (i.e., the same cell), even though they are two “topological objects” (i.e., two well separated connected components), we created the pair detection model to identify these pairs and correct the cell instances such that the pair of daughter cells have the same instance identity.

**Figure 14:**
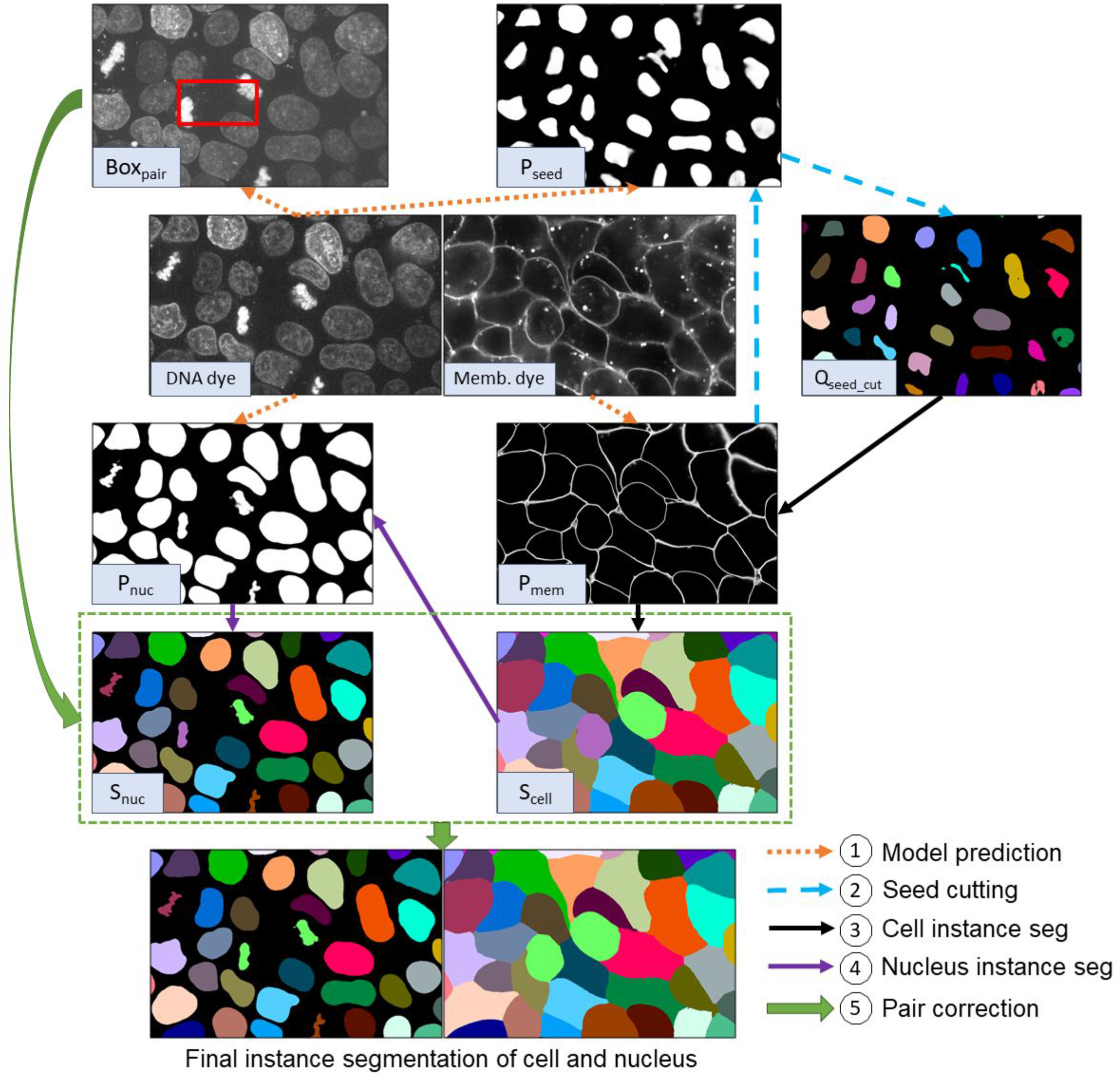
Illustration of the five key computational steps for obtaining the final instance segmentation of cells and nuclei from membrane dye and DNA dye images. The circled numbers and arrows in the workflow and the symbols in the bottom left box in each of the images represent the steps described in the method section.

### Label-free integration

When applying instance segmentation algorithms at scale, over many thousands of images, a particular source of variation that causes decreased segmentation performance is that the DNA and/or cell membrane dyes sometimes do not stain all cells equally and some cells quite poorly. Badly stained cells are extremely dim, or even barely visible, in the DNA and/or the cell membrane channels. Training on large sets of data does not improve the robustness in such scenarios due to the absence of these signals. In our case, this problem is very rare (less than 1% of cells) due to the rigorous quality controls of our imaging pipeline. However, when dealing with large numbers of cells even this small percentage represents many cells (e.g., 200,000 cells in total means 1% is 2000 cells). We can further avoid this type of error by integration of our segmentation workflow with the label-free method [Ounkomol et al., 2018]. Originally, the label-free method was designed to predict fluorescent images of different intracellular structures from bright field images in order to see different parts of a cell without any fluorescent marker. Here, we used the same idea to predict the segmentation mask from directly from bright-field images. Specifically, we trained the label-free model from bright-field images to cell boundary segmentations (using the output from the membrane dye based cell membrane segmentation model) and from bright-field images to nuclear segmentation (using the output from the DNA dye based nuclear segmentation model) on a selected set of images (∼400 images). In general, the label-free models by themselves cannot provide as accurate segmentation of cell and nuclei boundaries as the segmentation models using the fluorescent signals, but are still useful as “boosting” models to improve overall segmentation algorithm performance. The output from these two label-free models were combined with the semantic segmentation from membrane dye and DNA dye images to further improve the robustness. A representative example of the final “official names” segmentations is shown in **Fig. 17**, to demonstrate the overall quality of the cell and nuclear segmentation near the bottom, middle and top.

**Figure 17:**
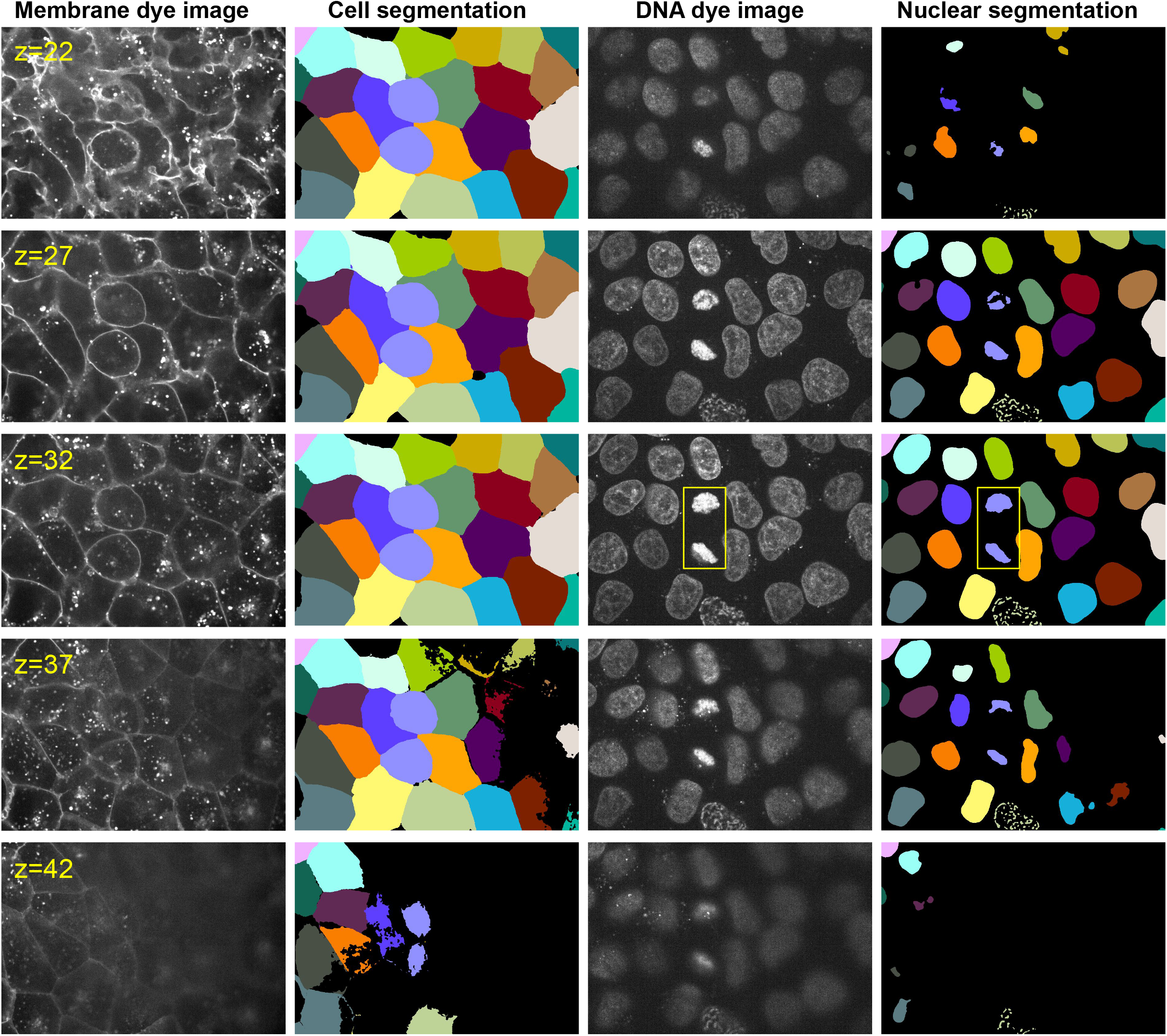
An overview of the final 3D instance segmentation of cells and nuclei from the DNA and membrane dye-based cell and nuclear segmentation algorithm. The results are shown at selected z-slices near the bottom, middle and top of the cells. The instances are represented by different colors. In z=22 and z=42, it seems like the nuclear segmentation has significant false negative errors, since we can still observe DNA dye signals in the image. This is due to the diffraction of light along z. The segmentations actually have reasonable accuracy in z, as we demonstrated in **Fig. 15**. The yellow boxes indicate one example of cells in telophase/cytokinesis. The pair correction step described in the Methods successfully pairs up the cells (both in purple color).

### Extending the lamin B1 segmentation “model” to a lamin B1 segmentation “algorithm” for more biologically accurate segmentation

In general, the models developed from the iterative deep learning workflows are neural networks that make per-voxel predictions of the probability of being part of the target structures. Even though the model can do a decent job in general, there is still no guarantee of the topological properties of the segmentation results, which is an important research area being actively studied in the computer vision community [Hu et al., 2019]. In the context of lamin B1 segmentation, one specific topological property that could be important for certain downstream analyses is the completeness of the segmented shell-shape representing the nuclear envelope. In other words, the interior of the segmented shell should be topologically disconnected from the exterior of the shell. Considering the nuclear envelope is very thin, a few pixels of false negative segmentation along the shell can alter this topology.

To solve this problem, we developed an algorithm that takes both the lamin B1 channel and the membrane dye channel as the input and utilizes four deep learning models including (1) the lamin B1 filled segmentation model we developed for nuclear segmentation training assay, (2) the overall lamin B1 segmentation developed with iterative deep learning workflow described in previous sections, (3) a lamin B1 seeding model, and (4) the membrane dye-based cell segmentation model. The overall steps are illustrated in **Fig. 18**. Specifically, with the output from the four models, we used the cell membrane segmentation model output (after binarization) to cut the lamin B1 seeding model outputs to generate one seed per interphase nucleus and also provide placeholder seeds for background objects. Then, the seeds were applied on the overall lamin B1 segmentation model output via a seeded watershed algorithm to obtain filled interphase nucleus segmentation. There may be false positive seeds due to the “bubble” part of lamin B1 in certain mitosis stages (see A and I in **Fig. 18**). Such seeds and the corresponding objects in the watershed output can be identified by comparing the watershed output with the lamin B1 fill segmentation model output. An object in the watershed output will be removed if it overlaps a lot with the background in the lamin B1 fill model output. After removing objects not corresponding to interphase nuclei, the boundaries of these objects are extracted and merged with the overall lamin B1 segmentation as the final lamin B1 segmentation result. The final result (see **K in Fig. 18**) contained both complete nuclear envelope and properly segment invaginations and lamin B1 during mitosis.

**Figure 18:**
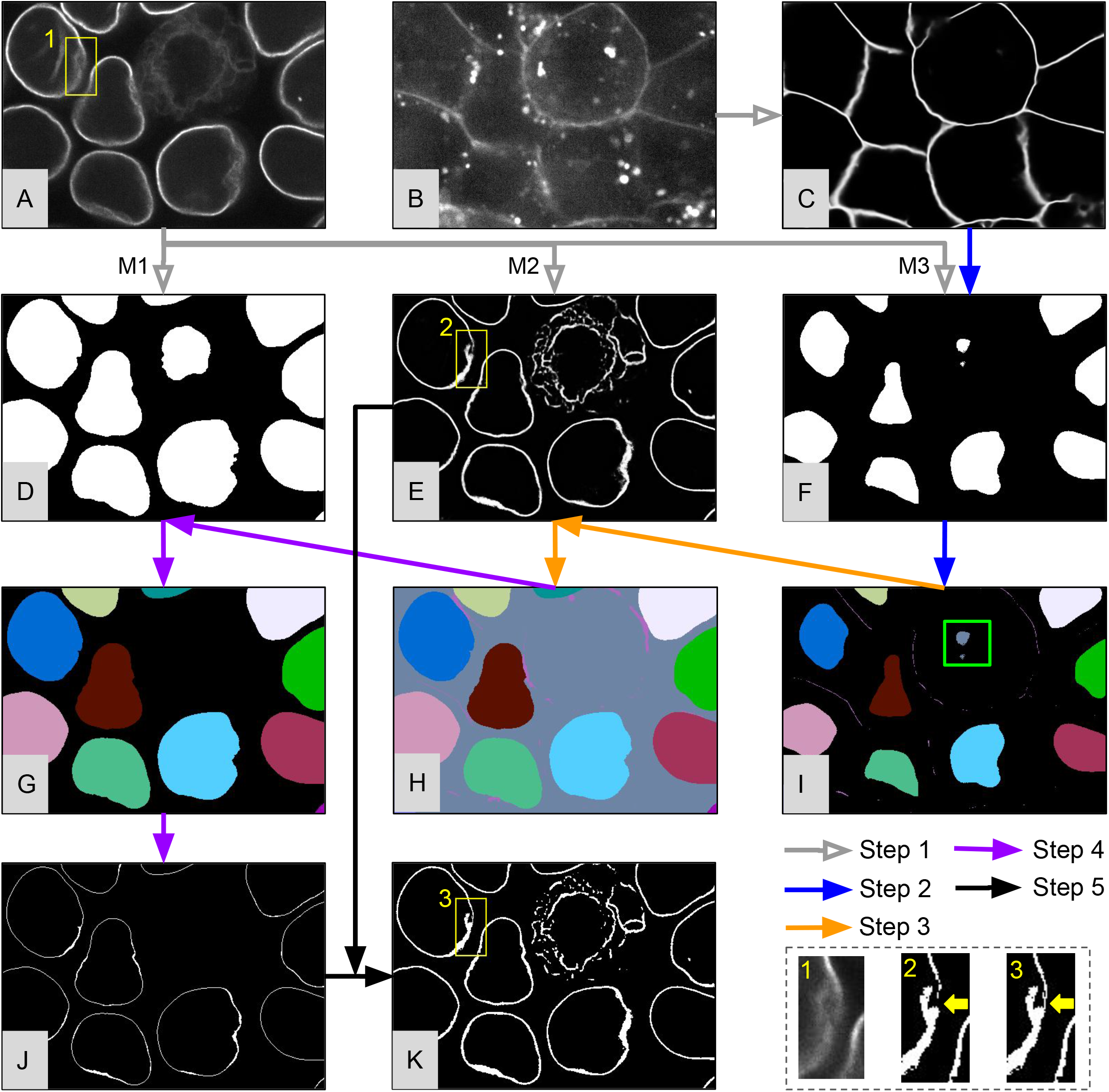
Illustration of the five key computational steps of the lamin B1 segmentation algorithm extended from the lamin B1 segmentation model. In step 1 (gray arrows), we apply the lamin B1 fill model (M1), the overall lamin B1 segmentation model (M2) created with the iterative deep learning workflow, and a lamin B1 seeding model (M3) to the lamin B1 image (see A), with results shown in D, E, F, respectively. We also apply the membrane dye-based cell segmentation model on the membrane dye image (see B) to obtain the cell boundaries (see C). In step 2 (blue arrow), a pruned seed image (see I) is created from C and F. The cell boundaries shown in C are used for two purposes: (1) to cut the seeding model output (F) to separate touching seeds, if any, and (2) to serve as extra seeds for background objects. The seed labeled in the green box in I is not necessary, as it corresponds to a mitotic cell, but will not cause errors in the final segmentation as the downstream steps are designed to be robust to this issue. In step 3 (orange arrows), a seeded watershed algorithm (with I as seeds) is applied on the overall lamin B1 segmentation model output (E) to obtain the interphase filled lamin B1 segmentation (see H). The background color of H is the same as the seed labeled in the green box in I, which indicates the false positive seed in I resulted in a background object. In step 4, we want to identify such falsely detected objects and remove them. We compare the lamin B1 fill model output (D) with the watershed output (H) to remove any large objects in H that overlap significantly with the background in D. The result is shown in G, and the boundary of each object is extracted as interphase lamin B1 shells (see J). Finally, the shells (J) are combined with the binarized overall lamin B1 segmentation model output (E) as the final result (see K). Three yellow boxes, marked by 1, 2, and 3, label a small area in A, E, and K. The zoomed-in images of these three boxes are shown in the lower right panel. We can see the result (see box 2) from the overall lamin B1 segmentation model from iterative deep learning workflows contains a broken shell due to the folding of the shell. The final version (see box 3) successfully generates a closed shell.

## Materials and Methods

### Data collection for toolkit development

The image segmentation toolkit has been applied to data produced at the Allen Institute for Cell Science using gene-edited, human induced pluripotent stem cells (hiPSCs) in both the undifferentiated stem cell and hiPSC-derived cardiomyocytes. Briefly, CRISPR/Cas9 was used to introduce mEGFP and mTagRFP-T tags to proteins localizing to known intracellular structures [Roberts et al., 2017, Haupt et al., 2018]. Clonal, FP-tagged lines were generated for each intracellular structure of interest and were used in imaging experiments in which undifferentiated hiPS cells were labeled with membrane dye (CellMask Deep Red) and DNA dye (NucBlue Live) to mark cell boundaries and the nucleus (see the SOP at allencell.org). Edited hiPSC cell lines were differentiated into cardiomyocytes using a small-molecule protocol, as described previously (allencell.org and [Roberts et al., 2018]). For imaging, cells were plated onto glass bottom plates coated with matrigel for undifferentiated hiPS cells and polyethyleneimine and laminin for cardiomyocytes (see SOPs at allencell.org), respectively and were imaged using a ZEISS spinning-disk microscope with a 100x/1.25 Objective C-Apochromat W Corr M27, a CSU-X1 Yokogawa spinning-disk head or a 40x/1.2 NA W C-Apochromat Korr UV Vis IR objective, and Hamamatsu Orca Flash 4.0 camera. Imaging settings were optimized for Nyquist sampling. Voxel sizes were 0.108 µm × 0.108 µm × 0.290 µm in x, y, and z, respectively, for 100x, hiPSC images and 0.128 µm × 0.128 µm × 0.290 µm in x, y, and z, respectively, for 40x, cardiomyocyte images. The mEGFP-tagged Tom20 line was transfected with mCherry-Mito-7 construct (Michael Davidson, addgene #55102) using 6 μl per well of transfection mixture containing 25 μl Opti-MEM (ThermoFisher #31985-070), 1.5 μl GeneJuice (Millipore #70967) and 1 ug endotoxin free plasmid. Transfected cells were imaged the next day on a ZEISS spinning disk confocal microscope as above. All channels were acquired at each z-step.

### Algorithms in the classic image segmentation workflow

#### Step 1: Pre-processing

To prepare images for segmentation, we first performed intensity normalization and smoothing. Our toolkit includes two normalization algorithms to choose from, min-max (MM) and auto-contrast normalization (AC). Min-max normalization transforms the full range of intensity values within the stack into the range [0,1]. Auto-contrast normalization adjusts the image contrast by suppressing extremely low/high intensities. To do this, the mean and standard deviation (std) of intensity is first estimated by fitting a Gaussian distribution to the whole stack intensity profile. Then, the full intensity range is cut off to the range [mean - a × std, mean + b × std], and then normalized to the range [0, 1] (**Fig. 3A**). The parameters, a and b, can be computed automatically based on a subset of typical images or can be user-defined. Auto-contrast is recommended by default. Min-max normalization should be used when the voxels with highest intensities are the target in the structure and should not be suppressed. For example, in “point-source” structures, such as centrin-2 (centrioles), the voxels with highest intensities usually reside in the center of the structure, making them critical to preserve. In addition to intensity normalization, there are three smoothing operations available in the pre-processing step: 3D Gaussian smoothing (G3), slice-by-slice 2D Gaussian smoothing (G2), and edge-preserving smoothing (ES; [Perona and Malik, 1990]). 3D Gaussian smoothing generally works well. However, if the target structure consists of dense filaments, an edge-preserving smoothing operation may be more effective (**Fig. 3B**). Slice-by-slice 2D Gaussian smoothing should be used when the movement of the intracellular structure is faster than the time interval between consecutive z-slices during live 3D imaging. In this situation, 3D smoothing may further aggravate the subtle shift of the structure in consecutive z-slices.

#### Step 2: Core segmentation algorithms

After pre-processing the images, they are segmented via a selection of core segmentation algorithms (**Fig. 1A**). 2D and 3D filament filters (F2 and F3; **Fig. 4A**; [Jerman et al., 2016]) are suitable for structures with curvi-linear shape in each 2D frame (e.g. Sec61 beta in **Fig. 4A**) or filamentous shape in 3D (e.g. alpha tubulin in **Fig. 4A**). The 2D and 3D spot filters (S2 and S3) compute the Laplacian of the Gaussian of the image in either 2D or 3D and thus can detect similar yet distinct spot-like localization patterns (**Fig. 4B**). The “point-source” desmoplakin localization pattern, exhibits as a round and fluorescence-filled shape in 3D. The S3 filter is more accurate for desmoplakin than the S2 filter, which stretches filled, round objects in the z-direction. For structures with a more general spotted appearance within each 2D frame instead of separate round structures (e.g. fibrillarin vs. desmoplakin in **Fig. 4B**), the S3 filter may fail to detect obvious structures while the S2 filter performs much better. The core watershed algorithm (W) can be used in two different ways. First, watershed can be applied to distance transformations of S3 filter results using local maxima as seeds to further separate proximal structures (**Fig. 4C**). Second, watershed can also be directly applied to the pre-processed image with seeds (detected by another algorithm) to segment structures enclosed in fluorescent shells (e.g. lamin B1 in **Fig. 4D**). The last core segmentation algorithm, masked object thresholding (MO) is designed for intracellular structure patterns with varying granularity or intensity (e.g. nucleophosmin in **Fig. 4E**). The MO threshold algorithm first applies an automated global threshold to generate a pre-segmentation result, which is used as a mask to permit an Otsu threshold to be applied within each pre-segmentation object. For example, the nucleophosmin localization pattern includes a primary localization to the granular component of the nucleolus and a weaker, secondary localization to other parts of both the nucleolus and nucleus. Therefore, we first apply a relatively low global threshold to roughly segment each nucleus We next compute a local threshold within individual nuclei to segment the nucleophosmin pattern (**Fig. 4E**). Compared to traditional global thresholding, masked-object thresholding performs more robustly to variations in intensity of the nucleophosmin localization pattern in different nuclei within the same image.

#### Step 3: Post-processing

Three different algorithms are available for the final post-processing step in the workflow (**Fig. 5**). These algorithms refine the binary core segmentation algorithm result to return a final segmentation. Not all post-processing algorithms are needed for every structure. The first algorithm is a morphological hole-filling algorithm (HF) that can resolve incorrect holes that may have appeared in certain segmented objects to represent the target structure more accurately (**Fig. 5A**). Second, a straightforward size filter (S) can be used to remove unreasonably small or large objects from the core segmentation algorithm result. (**Fig. 5A**). Finally, a specialized topology-preserving thinning operation (TT) can be applied to refine the preliminary segmentation without changing the topology (e.g., breaking any continuous but thin structures). This thinning is accomplished by first skeletonizing the preliminary segmentation, then eroding the segmentation in 3D on all voxels that are not themselves within a certain distance from the skeleton (**Fig. 5B**).

### Main computational steps in Training Assays for semantic segmentation of nuclei/mitotic DNA

**Fig. 12** is an overview of the eight key steps in a workflow to build a deep neural network for accurate semantic segmentation (entire field of view, not individually detected objects) of nuclei/mitotic DNA from DNA dye labeled cells via two Training Assays. Step (1) and (2) are the same as the first iteration in the iterative deep learning workflow for lamin B1 segmentation (see **Fig. 8**), except using the filled shell of lamin B1 segmentation as the segmentation target to train the “lamin B1 fill” model. (3) We applied this “lamin B1 fill” model to 1017 images to obtain the filled lamin B1 shells and use them as the segmentation target for interphase nuclei. (4) We took advantage of an earlier version of cell segmentation of the hiPSC Single-Cell Image Dataset, where the cell cycle stages were annotated, and trained a separate deep learning model to roughly estimate regions of mitotic DNA in these images to approximate the exclusion masks (i.e., the regions not considered during training) automatically. Alternatively, in the absence of pre-annotated mitotic cells, the exclusion of mitotic cells could have been performed with the Segmenter interface for the merging curator. Mitotic cells are 2∼5% of our cell population and therefore present about 50 images. So, performing this step manually would not have been overly time consuming. We used all filled nuclear segmentation images (excluding mitotic cells) as the ground truth to train a deep neural network model to segment interphase nuclei from DNA dye images (**DNA dye-based interphase nuclei segmentation model**). We chose to use a large number of training data images as DNA dye exhibits considerable image-to-image variation and we wanted to create a model that was sufficiently robust to be applied to over 18,000 FOV images of the hiPSC Single-Cell Image Dataset. Less training data would have sufficed for problems at a smaller scale. (5) Now that we had a successful model for interphase nuclei via DNA dye, we required an equivalently robust segmentation of mitotic DNA, which we could then use to merge with the interphase nuclei model to create a single DNA-dye based nuclear segmentation model. We turned to the mEGFP-H2B cell dataset and first applied the DNA dye-based interphase nuclei segmentation model to the DNA dye channel of these images to get an accurate segmentation of interphase nuclei. (6) In parallel, we developed classic image segmentation workflows in Segmenter specialized for mitotic DNA via mEGFP-H2B in prophase, metaphase, and anaphase (one workflow for prometaphase/metaphase, and one workflow for other mitosis stages). We did not include telophase/cytokinesis because that stage was already successfully segmented with the interphase nuclei model since the nuclear envelope was already reformed by this stage. We then applied these Segmenter workflows to the same mEGFP-H2B dataset as in step 5. (7) We then used Segmenter to perform a merging curation on the two segmentations from step 5 and step 6 to create a merged ground truth image set including both DNA-dye based interphase nuclei and mitotic DNA. (8) Finally, we trained a new model, the **DNA dye-based nuclear segmentation model** on this merged ground truth. This single model can now generate semantic segmentations of nuclei/mitotic DNA in all cell cycle stages from images of DNA dye labeled hiPSCs.

### Main computational steps for Training Assay for semantic segmentation of the cell membrane

**Fig. 13** is an overview of the four key steps in a workflow to build a deep neural network for accurate semantic segmentation of the cell boundary via a cell membrane dye image and a Training Assay. (1) We employed a semi-automatic algorithm based on a seeded watershed algorithm to obtain the initial semantic segmentation of the CAAX channel in the CAAX dataset (described just below). (2) We used the sorting curator in Segmenter to inspect the initial segmentation and selected 7 “passing” images from a pool of 10 images. A CAAX segmentation model was trained with the selected 7 images. Even though manual steps were involved in seed generation to guarantee the seeding quality for this first set of CAAX ground truth segmentations, the watershed algorithm still failed to segment certain cells due to the thin membrane at the top of the cells. The CAAX segmentation model can learn from the sorted successful examples to outperform the watershed algorithm (3) We applied the CAAX model to 312 images from the CAAX cell line to obtain the resultant successful CAAX segmentation of these images. (4) Then, we used these CAAX segmentations as the segmentation ground truth target to train **the membrane dye-based cell segmentation model** that successfully segments cell boundaries from images of membrane dye stained hiPSCs images.

To create the initial CAAX segmentation, we used a seeded watershed algorithm. The seeds were obtained in a semi-automatic manner. First, we applied the cell seeding model, described in detail in the Seeding model section just below, to generate initial seeds (roughly one connected component centered inside each nucleus). Then, we manually corrected the initial seeds by (a) cutting the falsely merged seeds, (b) adding one seed manually (e.g., painting with one stroke in ImageJ) inside each nucleus of any cell that was only partly within the field-of-view and thus lacked automatically predicted seeds via the seeding model, (c) manually extending some seeds (less than 10%; e.g., painting in ImageJ with initial seeds overlaid on CAAX image), where the cells had elongated extensions that we needed to ensure the seeds were able to grow into by the watershed algorithm, (d) adding an auxiliary bottom seed on the bottom slice and an auxiliary top seed on the top slice as “placeholders” for the objects representing background in the bottom and the top.

We upsampled the 3D microscopy image to isotropic (from 0.108 x 0.108 x 0.29 um to 0.108 x 0.108 x 0.108 um) with cubic interpolation and upsampled the seed image accordingly with no interpolation. Then, we performed a 3D Gaussian smoothing on the upsampled CAAX image and performed a seeded watershed on the smoothed image with the seeds that were semi-automatically generated above. In general, the watershed algorithm with no watershed line is more suitable for detecting the thin membrane at the top of cells, while the watershed algorithm with watershed line achieves more accurate separation boundaries between cells. We therefore ran the watershed algorithm twice, once with and once without watershed line, both with a connectivity of 26 in 3D. The results were denoted as W_line_ and W_no_line_.

Finally, we want to merge W_line_ and W_no_line_ into a CAAX watershed-based segmentation workflow. Roughly speaking, W_line_ and W_no_line_ are merged based on the watershed object grown from the top auxiliary seed in W_no_line_ (denoted by C_top_). Let B_1_ be the watershed line in W_line_, which is equivalent to the cell membrane except the sub-optimal membrane at the top of cells. Let B_2_ be the boundaries of all cells in W_no_line_, which creates a better estimation of the membrane at the top of cells than B_1_. The final CAAX segmentation workflow is B_1_ with voxels replaced by B_2_ if less than 3 voxels away from C_top_.

### Cell seeding model for instance segmentation of cells and nuclei

The goal of the cell seeding model is to predict one single connected component for each individual cell from DNA dye images. The seeding model is similar to the DNA dye-based nuclear segmentation model but predicts a slightly different mask. For cells in interphase, we created the seed mask as a “shrinked” nuclear segmentation mask. This was implemented as a morphological erosion of the nuclear segmentation mask with a disk-shape structure element of radius 15, then keeping only the largest connected component in the eroded result. This “shrink mask” aims to encourage the seeding model to predict two separated seeds when two nuclei are tightly touching each other. For cells in mitosis, we created seed masks to be the convex hull of the mitotic DNA segmentation mask for each cell. However, mitotic cells in later anaphase and telophase/cytokinesis required different treatment because the mitotic DNA was now separated into two regions of the cell or into the two daughter nuclei reforming in the daughter cells. These specific cells are permitted to, and in fact should have, two seeds per cell, one for each of the mitotic DNA/reforming nuclei that are pulled apart within a single cell. So, when creating the seed mask for these cells, two convex hulls are computed, one for each of the mitotic DNA/reforming nuclei, instead of just one for the whole cell. We used the merging curator in Segmenter to create a merged ground truth image set of 600 images consisting of the interphase and mitotic seed masks. This new merged ground truth set was used to train the final cell seeding model that was then used to create the seeds for the CAAX watershed segmentation results.

### Pair detection model for refining instance segmentation of cells and nuclei

We need each cell to have exactly one predicted seed from the seeding model in order to be able to perform the Watershed to create a CAAX-based ground truth image set as a Training Assay. But, as we discussed in the seeding model section above, cells in late anaphase or telophase/cytokinesis naturally have two predicted seeds for each of the mitotic DNA/reforming nuclei that are pulled apart within a single cell. To address this issue, we trained a pair detector based on FasterRCNN [Ren et al., 2015]. The detector can automatically find pairs of mitotic DNA/reforming nuclei belonging to the same cell in late anaphase or telophase/cytokinesis and return a bounding box tightly encompassing each pair of mitotic DNA/reforming nuclei in late anaphase or telophase/cytokinesis. Such bounding boxes are then used to locate the nuclei pairs.

We applied the cell and nucleus segmentation algorithm (without pair correction) on a manually selected subset of 60 images from the hiPSC Single-Cell image Dataset, where each image has at least one pair of cells in late anaphase or telophase/cytokinesis. We created maximum intensity projections along z of the segmentation results and calculate a 2D bounding box for each mitotic DNA pair based on their segmentation. We then used these bounding boxes as the ground truth to train a 2D Faster-RCNN model to detect mitotic DNA pairs from the maximum intensity projections of the DNA dye image. We used the off-the-shelf implementation from https://github.com/facebookresearch/maskrcnn-benchmark. The model was trained with learning rate 0.001. We validated the performance of this model with a set of held-out images with about 30 cell pairs in total. The model made 0 false negative errors, which is critical to the success of cell pairs in the final segmentation, since missed detection cannot be recovered. We applied the detection model on our large dataset and spot-checked the performance on selective sample data to ensure the performance remained consistent with prior validation.

### Main computational steps for instance segmentation of cells and nuclei

The prediction outputs from the nucleus/mitotic DNA segmentation model, cell membrane segmentation model, seeding model and the pair detection model are all at the level of the entire FOV and thus are semantic segmentation predictions. We combined these models and designed a DNA and membrane dye-based cell and nuclear segmentation algorithm to generate instance segmentations that identifies all individual cells and nuclei/mitotic DNA in an image. The algorithm required five key steps as depicted in **Fig. 14**. (1) Model prediction: We applied the DNA dye-based nuclear segmentation model, the seeding model, and the pair detection model on the DNA dye channel. The predictions of these three models are denoted by P_nuc_, P_seed_ and Box_pair_, respectively. We applied the membrane dye-based cell membrane model on the membrane dye channel. The prediction is denoted by P_mem_. (2) Seed cutting: P_mem_ and P_seed_ are first binarized (by a cutoff value determined from sample data) into Q_mem_ and Q_seed_, respectively. Then, foreground voxels in Q_mem_ are removed from the foreground of Q_seed_ and the result is called Q_seed_cut_. Minor refinement is performed on Q_seed_cut_, such as removing small objects (see released code for details). The final seed image Q_seed_final_ is Q_seed_cut_ with one extra seed for the stack bottom and one extra seed for the stack top as axillary seeds as described in Watershed section above. (3) Cell instance segmentation: Perform seeded watershed on P_mem_ with Q_seed_final_ as seeds. The objects grown from the axillary seeds are ignored (identified by object indices). The top and bottom axillary seeds were used as “placeholder” objects to represent background near the top and bottom of the image. The bottoms of the cells interact with the substrate and cells form many protrusions that can overlap each other, while the z-resolution does not allow proper disentangle the overlapping parts. All together the true boundary of the very bottom of the cell right at the substrate is very difficult to identify even by eye. Therefore, we decided to find a z slice near the bottom of the cell before the cell boundary becomes obscure and propagate that particular cell boundaries downward to the bottom of the cell. One example of the cell segmentation near the bottom of cells is presented in **Fig. 19**. To achieve this goal, the segmentation at the bottom of cells is refined by finding a proper z-slice, say Z_0_, near the bottom and replace the segmentation from the bottom z-slice of all cells to Z_0_ with segmentation in Z_0_ (see released code for details). The final cell instance segmentation result is denoted by S_cell_. (4) Nucleus instance segmentation: P_nuc_ is binarized (by a cutoff value determined from sample data) into Q_nuc_. Then, the cell instance labels from S_cell_ are propagated to Q_nuc_.

**Figure 19:**
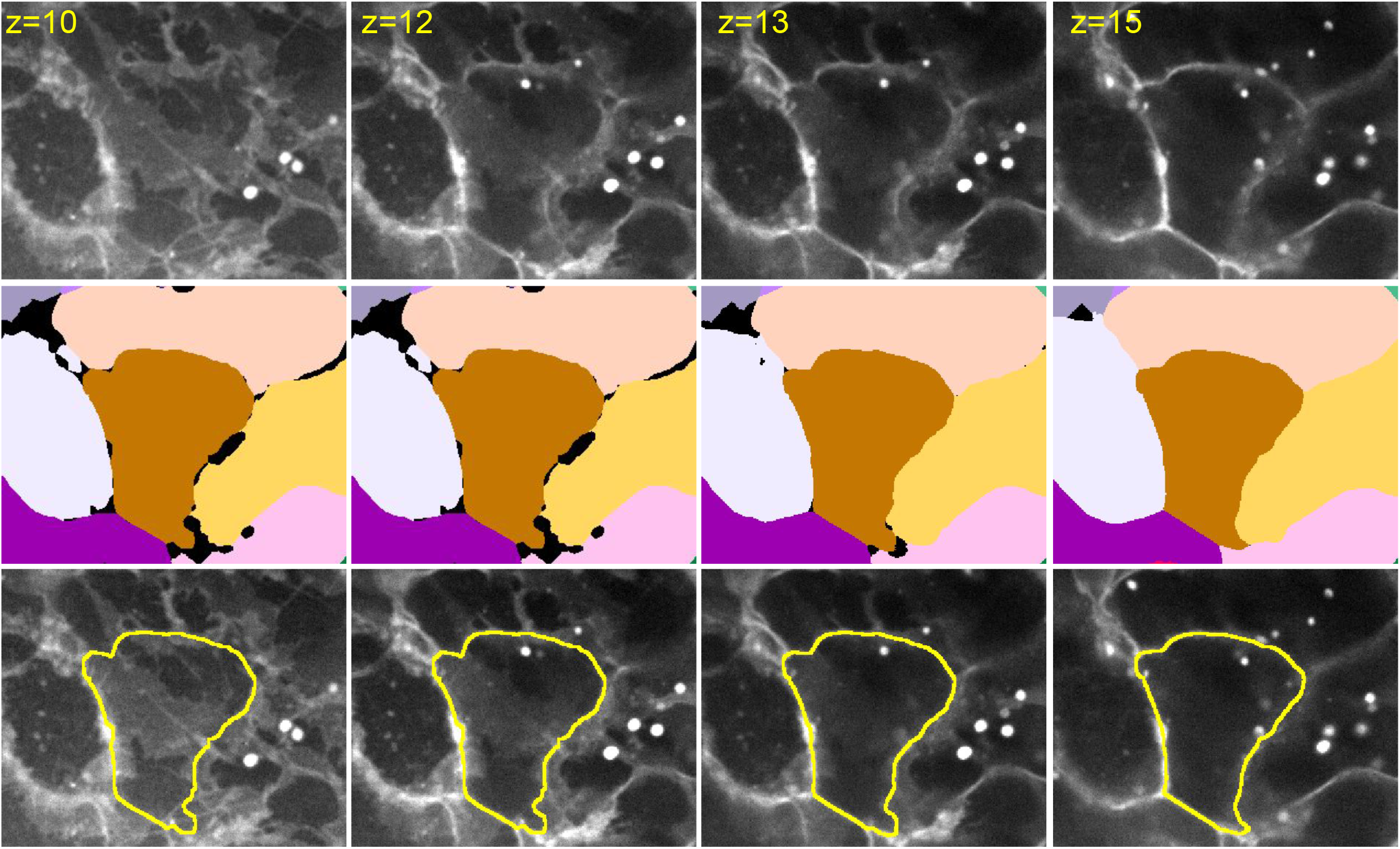
Examples of cell segmentation from the 3D cells and nuclei instance segmentation algorithm near the bottom of cells at selected z-slices. Rows from top to bottom: cell membrane dye images, cell instance segmentation, cell membrane dye images overlaid with the boundary of the cell near the center. Segmentation in z=12 is automatically picked as a reasonable segmentation near the bottom and is duplicated down to z=10, in which the boundary of each cell is difficult to tell.

The result after slight refinement (e.g., filling holes, see released code for details) is denoted as S_nuc._ (5) Pair correction: All detected bounding boxes from Box_pair_ are overlaid onto S_nuc_ to extract the instance indices of each mitotic cell pair. If only one instance or nothing exists in the bounding box, this box is treated as a false positive and ignored; otherwise, the mitotic cell pairs are verified by checking the confidence score from the pair detection model, the relative intensity and size of the two parts of the cell, and how much of each of the two objects is enclosed by the bounding box (see details in the released code). For each confirmed pair, the corresponding two instance indices are merged to the same index.

### Computational steps for integrating label free for instance segmentation of cells and nuclei

We trained two label-free models to predict cell boundary segmentation and nuclear segmentation directly from bright-field images. Suppose the two label-free models are LF_nuc_ and LF_mem_. These two models can be used to boost the robustness of final cell and nuclear instance segmentation, even though they may not be accurate enough by themselves. Specifically, the algorithm for instance segmentation of cells and nuclei above can be slightly modified as follows to integrate LF_nuc_ and LF_mem_. (1) The output of LF_mem_ was combined with P_mem_ by weighted sum and this result was used as the cell boundary prediction. (2) The output of LF_nuc_ was first binarized by an empirically determined threshold (0.25) and then processed similarly to the seed cutting process described above. In the result image, any connected component that is not in Q_seed_ or Q_nuc_ will be added to Q_seed_ or Q_nuc_, respectively.

### Evaluation of cell and nuclear 3D instance segmentations

There is no “absolute ground truth” for us to quantitatively evaluate the quality of our instance segmentation of cells and nuclei. We therefore used an evaluation strategy combining a voxel level qualitative assessment and an instance level quantitative assessment. To evaluate the quality of nuclear segmentation at the voxel level, we use another specialized image-based biological assay, cells expressing mEGFP-tagged nucleoporin Nup153. Like lamin B1, nucleoporin Nup153 forms a “shell” around the nucleus, as nucleoporin Nup153 is located on the nuclear side of the nuclear pore complex. Nucleoporin Nup153 tags the nuclear pores and thus appears like densely packed puncta along the surface of the nucleus, instead of a continuous shell like lamin B1. Puncta along the surface of the nucleus suffer even less from the diffraction of light compared to even the continuous shell of lamin B1 and thus nucleoporin Nup153 data serves the purpose of evaluating the accuracy of the segmented nuclear boundary very well. If the nucleus segmentation is accurate, the boundary of the segmentation should reside near the center of those puncta along the nuclear boundary via tagged Nup153 in 3D. **Fig. 15** shows a visualization of this qualitative nuclear segmentation accuracy at the voxel level, which was very accurate.

**Figure 15:**
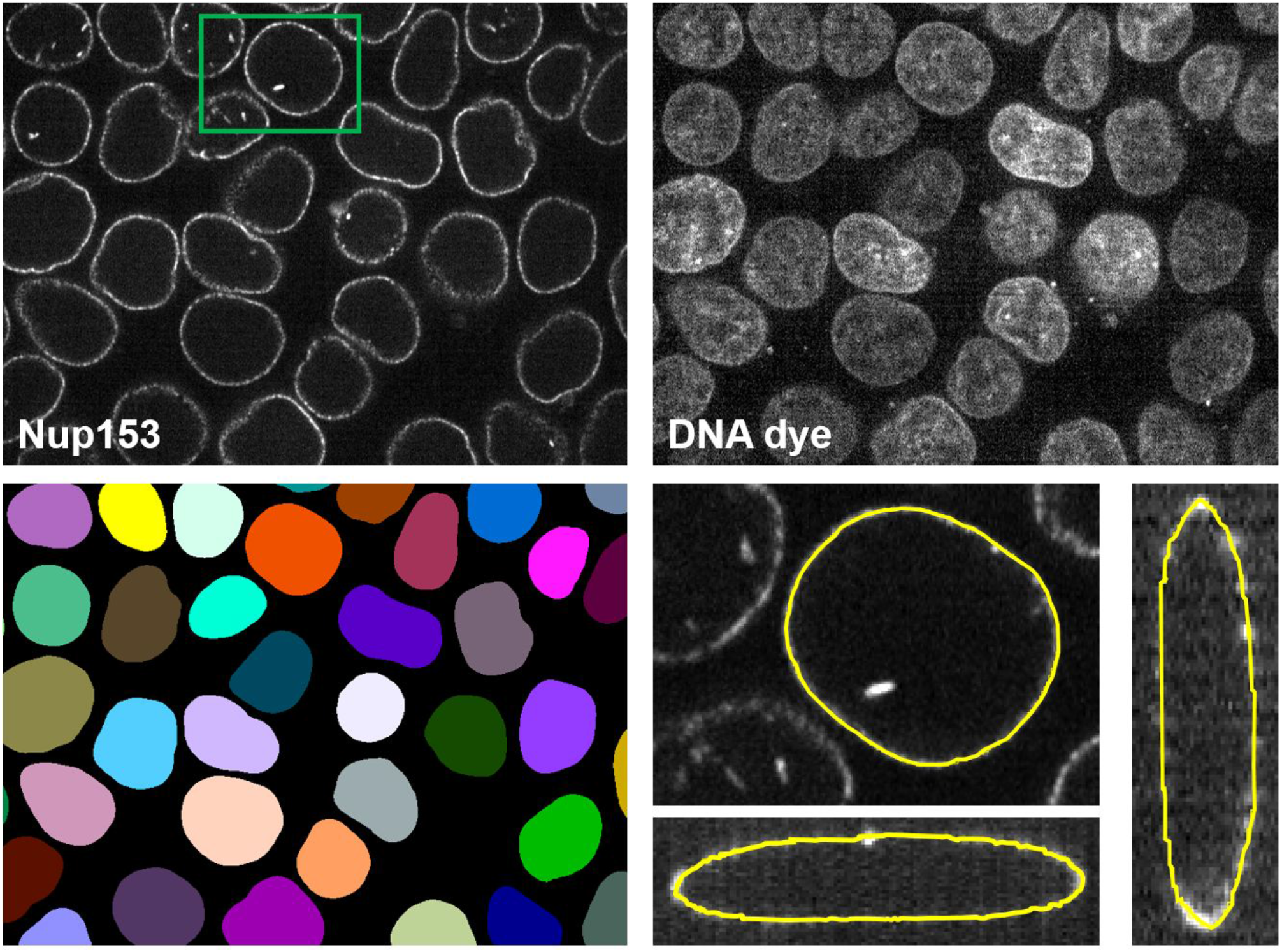
Qualitative evaluation of nuclear segmentation accuracy in the instance segmentation of cells and nuclei at the voxel level using mEGFP-tagged nucleoporin Nup153 images as a validation dataset. Nup 153 labels the nuclear pores and forms a shell of puncta around the nucleus boundary. Top row: a multi-channel 3D microscopy image (showing a single z slice) with both DNA dye (top right panel) and nucleoporin Nup153 (top left panel) of the same field-of-view. Bottom left panel: The instance segmentation of all nuclei from the DNA dye image (the result of the 3D cell and nuclei instance segmentation algorithm). Bottom right panel: Zoomed-in images of the nucleus in the green box in the top left panel, with front view and orthogonal views. The segmented boundary (the yellow lines) extracted from the segmentation result is overlaid on the zoomed-in images. The segmented nuclear boundary lies approximately at the center of the “shell” formed by the tagged nucleoporin Nup153 signal, which confirms the biological correctness of the instance segmentation of the nuclei via DNA dye and the generalizability of the DNA dye-based nuclear segmentation model, as this result is based on an entirely different dataset than the lamin B1 dataset the model was trained on.

To qualitatively evaluate the performance of the cell instance segmentation at the voxel level, we examined a small subset of the CAAX dataset held-out during model training. A sample result is shown in **Fig. 16**. Comparison on the front view (XY) of the segmented cells and the CAAX image shows mostly accurate cell boundaries even under considerable “noise” (due to the membrane dye labeling also labeling endocytic vesicles that were internalized during and after the membrane staining procedure) and correct cell instance identification, as best determinable by human experts. The comparison between the segmented cell boundaries and the CAAX image via the lateral side views (in z direction) can demonstrate the effectiveness of the membrane dye-based cell boundary segmentation model in achieving accurate segmentation of the membrane at the top of cells, even when parts of the membrane top signal from the membrane dye are low in brightness and contrast, contain artifacts from the dye, etc. We also observed that the segmented cell boundary mostly follows along the center of the cell membrane signal marked by CAAX, which confirms the biological correctness of the cell segmentation.

**Figure 16:**
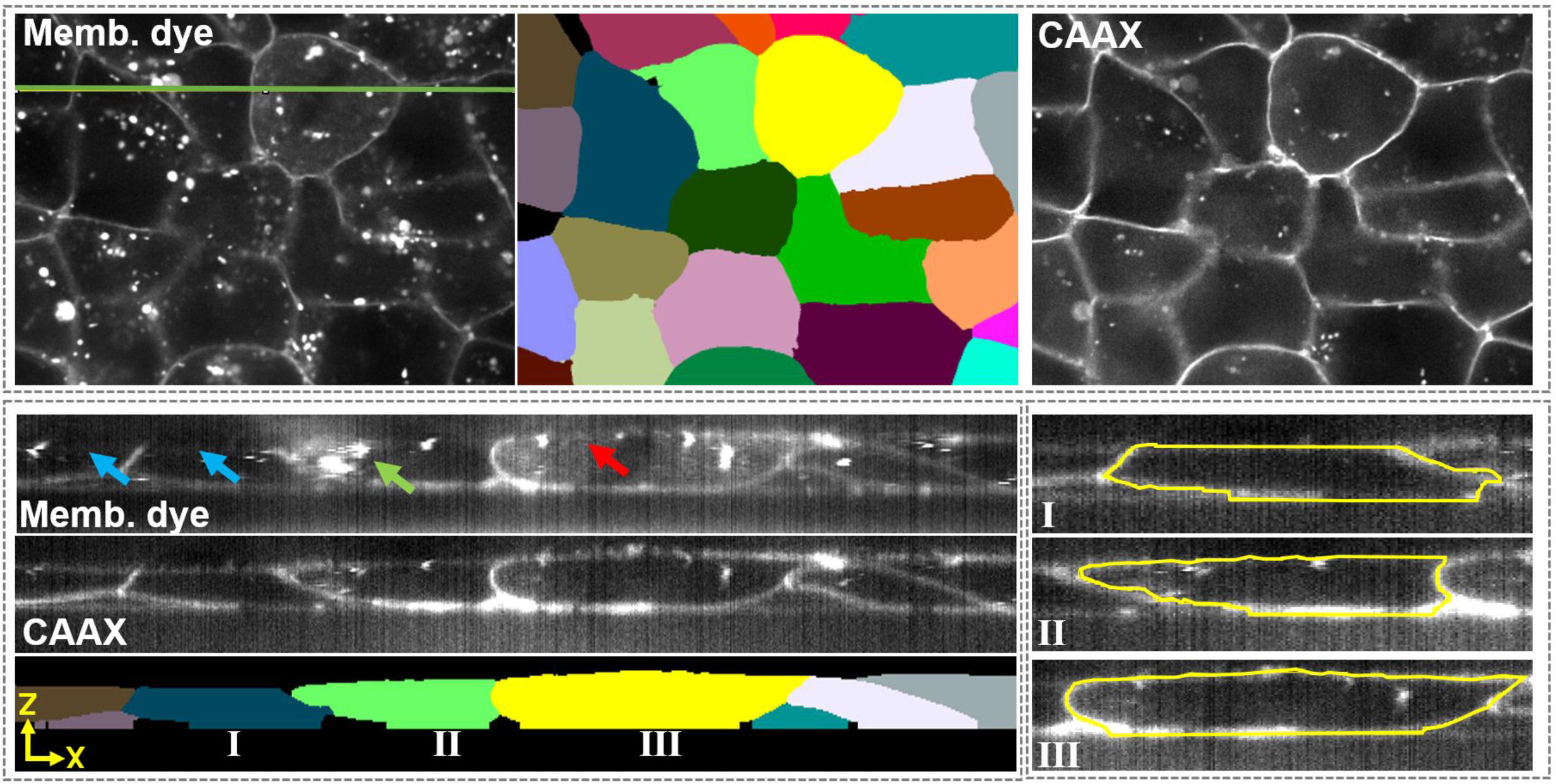
Qualitative evaluation of membrane dye-based cell segmentation using a subset of CAAX images held out from the model training. The three images in the top panel show the membrane dye (labeled “Memb. dye”), cell instance segmentation (center top panel) and CAAX images to verify the overall accuracy in XY. The lower left panel shows side views along the green line in the membrane dye image in the top panel to demonstrate the effectiveness of the model in achieving accurate detection of the membrane even with the low SNR of the membrane dye image near the top of the cells. The blue arrows indicate some areas with especially dim membrane signal. The red arrow indicates an area with very low contrast, but not very dim signal. The green arrow indicates an area where the membrane dye is significantly affected by the dye labeling endocytic vesicles. The bottom right panel shows zoomed-in images of three cells marked by I, II, III in the bottom image in the bottom left panel. The segmented boundary extracted from the cell instance segmentation is overlaid onto the zoomed-in CAAX images. We can observe that the segmented boundary lies approximately along the center of the cell membrane marked by CAAX, which confirms the biological validity of the cell segmentation.

After repeating the above qualitative inspection of cell and nuclear segmentation at the voxel level on about 20 randomly selected images, we confirmed the segmented cells and nuclei demonstrated sufficient accuracy to justify application of this workflow on a larger part of the dataset to extract all individual cells and nuclei in each field of view. We next evaluated the robustness of this segmentation accuracy over larger sized image sets at the instance level. We assessed a set of 576 images from 22 different cell structure lines to determine the percentage of entire FOV’s that contained successful cell and nuclear segmentation in 3D via the DNA dye and membrane dye. Each image was manually scored by at least two human experts using an in-house scoring interface. We found that over 98% of individual cells had successful cell and nuclear segmentation and over 80% of all images having the entire FOV successfully segmented.

### Deep Learning Models and Training

The deep learning models employed in our iterative deep leaning workflow are two fully convolutional networks specially customized for 3D fluorescence microscopy images: Net_basic and Net_zoom (**Fig. 20**; [Long et al., 2015] for more details about fully convolutional networks). Net_basic is a variant of a 3D U-Net [Çiçek et al., 2016] with (1) max pooling in all xyz dimensions replaced by max pooling in xy only, (2) zeros padding removed from all 3D convolution and (3) auxiliary loss added for deep supervision [Chen et al., 2016A]. Net_zoom has a similar architecture to Net_basic, but with an extra pooling layer with variable ratio to further enlarge the effective receptive field. Such modifications are made to deal with anisotropic dimensions common in 3D microscopy images and to improve the performance in segmenting tenuous structures, such as the thin nuclear envelope.

**Figure 20:**
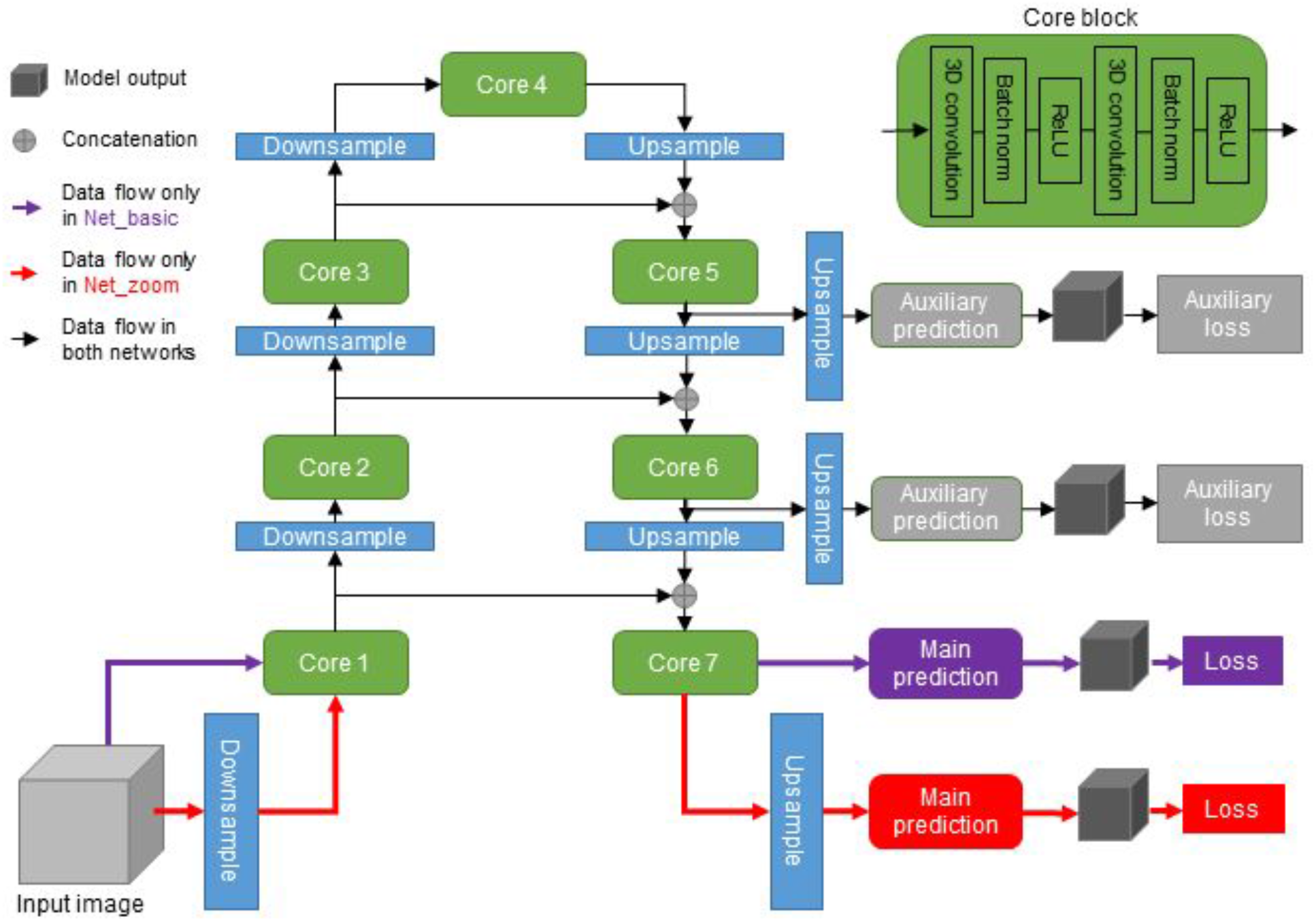
The architecture of the two deep neural networks used in the iterative deep learning workflow. The two networks, Net_basic and Net_zoom are almost identical in architecture. The layers and data flows that differ between Net_basic and Net_zoom are marked in purple and red, respectively. In general, the network consists of 7 core blocks connected by downsampling and upsampling layers. All core blocks have the same layers, detailed in the corner green box (two consecutive sets of 3D convolution with kernel size 3, batch normalization and ReLU activation). Both networks are attached to one main prediction branch and two auxiliary prediction branches. The main prediction block is one 3D convolution with kernel size 1, while auxiliary blocks have one 3D convolution with kernel size 3, followed by another 3D convolution with kernel size 1.

In each training iteration, random data augmentation is applied on each image and a batch of sample patches are randomly cropped from the augmented images. In practice, the patch size (i.e., the size of model input) and batch size (i.e., the number of samples trained simultaneously in each iteration) depend on the available GPU memory. For example, a single Nvidia GeForce GPU with 12GB memory is used in our experiments. With this setup, we choose a batch size of 4 and each input sample patch has size 140 × 140 × 44 voxels for Net_basic and 420 × 420 × 72 voxels for Net_zoom. For data augmentation, we adopt a random rotation by θ (a random value from 0 to π) and a random horizontal flip with probability 0.5. Weighted cross-entropy is used in all the loss functions, where a per-voxel weight is taken as a separate input image (e.g., cost map). By default, we use a weight = 1 for all voxels, but one can assign a larger weight on those extremely critical regions or assign zeros to those regions that do not count for the loss function. Models are trained with Adam [Kingma and Ba, 2014] with constant learning rate 0.00001 and L2 regularization with weight 0.005. For the nuclear segmentation model, cell membrane segmentation model, and the seeding model, we used the Net_zoom model (with zoom ratio 3).

### Segmentation algorithm comparisons

Representative sample images from 20 structure localization patterns were used to compare segmentation results between the Segmenter classic image segmentation workflow, 14 global and 8 local thresholding algorithms (**Fig. 7**). All images corresponded to the maximum intensity projection of a z-slice of choice plus and minus one z-slice. Each image was segmented with 14 automatic global thresholding algorithms available in ImageJ under the Image/Adjust/Auto Threshold option and 8 local thresholding algorithms available in ImageJ under the Image/Adjust/Auto Local Threshold option. The local algorithms require an additional parameter that corresponds to the radius of the local domain over which the threshold will be computed. Radius values 8, 16, 32 and 64 were tested for each local thresholding algorithms. For each of these results, the segmentation algorithms that provided the segmentation most similar to the Segmenter result was identified based on the Dice Metric. Global threshold algorithms included Huang, Intermodes, IsoData, Li, MaxEntropy, Mean, Minimum, Moments, Otsu, Percentile, RenyiEntropy, Shanbhag, Triangle, Yen. Local threshold algorithms included Bernsen, Contrast, Mean, MidGrey, Niblack, Otsu, Phansalkar, Sauvola.

### Determining mitochondrial width from published EM images

## Methods

Mitochondrial widths were determined in human pluripotent stem cells and human embryonic stem cells using previously published EM images [Bukowiecki et al. 2014, Niclis et al. 2015], respectively. JPEG versions of the EM images obtained from the manuscripts were opened in Fiji and mitochondrial width was measured for 5-10 mitochondria per EM image. A line was manually drawn between the outer mitochondrial membranes along the smaller mitochondrial axis. Line lengths were measured and converted into nanometers using the original scale bars in the figures. Mitochondrial width was found to be 256 +/- 22 nm for human pluripotent stem cells and 265 +/- 34 nm for human embryonic stem cells (mean +/- 95% confidence interval). An average mitochondrial width of 260 nm was used in **Fig 6**.

## Acknowledgements

We thank the entire Allen Institute for Cell Science team, who generated and characterized the gene-edited hiPS cell lines, developed image-based assays, recorded the high-replicate data sets suitable for image processing method development, and created the software infrastructure without whom this work would not have been possible. We especially thank the Allen Institute for Cell Science Gene Editing, Assay Development, Microscopy, and Pipeline teams for providing cell lines and images. We thank Jamie L. Gehring, Sara Carlson, Antoine Borensztejn for manually scoring the cell and nuclear segmentation results for evaluation. We thank Winfried Wiegraebe, Derek Thirstrup, Rick Horwitz, Gaudenz Danuser, Wallace Marshall, Tom Misteli, Jennifer Lippincott-Schwartz, and the Allen Institute for Cell Science Assay Development team for many insightful discussions. We also thank Matt Bowden and Basudev Chauduri for technical support related to software package release. We thank the Gladstone Institute for providing initial wild-type stem cell lines. We thank the Allen Institute for Cell Science founder, Paul G. Allen, for his vision, encouragement, and support.

