## supplemental information for "The Allen Cell and Structure Segmenter: a new open source toolkit for segmenting 3D intracellular structures in fluorescence microscopy images"

**1. Pseudocode of classic image segmentation workflow for sialyltransferase 1**

```
Input: I (original single channel 3D image stack)

Output: Final_segmentation (binary image of segmentation result)

Constant Parameters:

    normalization_param = [9, 19]

    G3_param = 1

    MO_param = [`tri`, 1200, False, True]

    S3_param = [[1.6, 0.02]]

    thin_param = [1, 1.6]

    min_size = 10

# pre-processing

I_norm = Auto_Contrast(I, normalization_param)

I_smooth = Gaussian_Smoothing_3D(I_norm, G3_param)

# apply S3 filter

PreSeg1 = Spot3D(I_smooth, S3_param)

# apply Masked Object Thresholding (MO) and thinning

PreSeg2 = MO_Thresholding(I_smooth, MO_param)

PreSeg2_thin = Topology_Preserving_Thinning(PreSeg2, thin_param)

# combine the results

Seg = Logical_OR(PreSeg1, PreSeg2_thin)

# size filtering

Final_segmentation = Size_Filter(Seg, min_size)
```

### 2. Pseudocode of classic image segmentation workflow for fibrillarin

```
Input: I (original single channel 3D image stack)
Output: Final_segmentation (binary image of segmentation result)
Constant Parameters:
    normalization_param = [0.5, 18]
    G3_param = [1]
    S2_param = [[1,0.01]]
    min_size = 5
# pre-processing
I_norm = Auto_Contrast(I, normalization_param)
I_smooth = Gaussian_Smoothing_3D(I_norm, G3_param)
# apply S2 filter
Seg = Spot2D(I_smooth, S2_param)
# size filtering
Final_segmentation = Size_Filter(Seg, min_size)
```

### 3. Pseudocode of classic image segmentation workflow for nucleophosmin

```
Input: I (original single channel 3D image stack)
Output: Final_segmentation (binary image of segmentation result)
Constant Parameters:
    normalization_param = [0.5, 15]
    G3_param = 1
    MO_param = [`ave`, 700, True, True]
    S2_param = [[2, 0.025]]
    S2_dark_param = [[2, 0.025], [1, 0.025]]
    min_size = 5
# pre-processing
I_norm = Auto_Contrast(I, normalization_param)
I_smooth = Gaussian_Smoothing_3D(I_norm, G3_param)
```

```
# apply Masked Object Thresholding (MO)
PreSeg1, MO_Mask = MO_Thresholding(I_smooth, MO_param)
# apply S2 filter to detect extra spots
ExtraSpots = Spot2D(I_smooth, S2_param)
PreSeg2 = ExtraSpots within MO_Mask
# apply S2 filter to detect dark spots
DarkSpots = Spot2D(1 - I_smooth, S2_dark_param)
# combine the results
Seg = Logical_OR(PreSeg1, PreSeg2) - DarkSpots
# size filtering
Final_segmentation = Size_Filter(Seg, min_size)
```

### 4. Pseudocode of classic image segmentation workflow for Sec61 beta

```
Input: I (original single channel 3D image stack)
Output: Final_segmentation (binary image of segmentation result)
Constant Parameters:
    normalization_param = [2.5, 7.5]
    F2_param = [[1,0.15]]
    min_size = 15
# pre-processing
I_norm = Auto_Contrast(I, normalization_param)
I_smooth = Edge_Preserving_Smoothing(I_norm)
# apply F2 filter
Seg = Filament2D(I_smooth, F3_param)
# size filtering
Final_segmentation = Size_Filter(Seg, min_size)
```

### 5. Pseudocode of classic image segmentation workflow for tom20

```
Input: I (original single channel 3D image stack)
Output: Final_segmentation (binary image of segmentation result)
```

**Constant Parameters:**

normalization_param = [3.5, 15]

G3_param = [1]

F2_param = [[1.5,0.16]]

min_size = 10

***# pre-processing***

*I_norm* = **Auto_Contrast**(*I,* normalization_param)

*I_smooth* = **Gaussian_Smoothing_3D**(*I_norm, G3_param*)

***# apply F2 filter***

*Seg* = **Filament2D**(*I_smooth*, F2_param)

***# size filtering***

*Final_segmentation* = **Size_Filter**(*Seg,* min_size)

### 6. Pseudocode of classic image segmentation workflow for LAMP-1

**Input:** *I* (original single channel 3D image stack)

**Output:** *Final_segmentation* (binary image of segmentation result)

**Constant Parameters:**

normalization_param = [3, 19]

G2_param = 1

S2_param = [[5,0.09], [2.5,0.07], [1,0.01]]

F2_param = [[1,0.15]]

hole_param = [1600, True]

min_size = 15

***# pre-processing***

*I_norm* = **Auto_Contrast**(*I,* normalization_param)

*I_smooth* = **Gaussian_Smoothing_2D_slice_by_slice**(*I_norm,* G2_param)

***# apply S2 filter***

*PreSeg1* = **Spot2D**(*I_smooth*, S2_param)

***# apply F2 filter***

*PreSeg2* = **Filament2D**(*I_smooth*, F2_param)

```
# combine the results
Seg = Logical_OR(PreSeg1, PreSeg2)
# hole Filling
Filled_Seg = Hole_Filling(Seg, hole_param)
# size filtering
Final_segmentation = Size_Filter(Filled_Seg, min_size)
```

**7. Pseudocode of classic image segmentation workflow for centrin-2**

```
Input: I (original single channel 3D image stack)
Output: Final_segmentation (binary image of segmentation result)
Constant Parameters:
    normalization_param = [8000]
    G2_param = [1]
    S3_param = [[1,0.04]]
    min_size = 3
# pre-processing
I_norm = Auto_Contrast(I, normalization_param)
I_smooth = Gaussian_Smoothing_2D_slice_by_slice(I_norm, G2_param)
# apply S3 filter
PreSeg = Spot3D(I_smooth, S3_param)
# watershed to cut falsely merged dots
Mask = Size_Filter(PreSeg, min_size)
Seed = Dilation( Local_maximum(I_norm), structure_element =
       ball_radius_1)
Watershed_map = -1 * Euclidean_distance_transform(Mask)
Watershed_seg = Watershed(Watershed_map, Seed, Mask)
# size filtering
Final_segmentation = Size_Filter(Watershed_seg>0, min_size)
```

### 8. Pseudocode of classic image segmentation workflow for desmoplakin

```
Input: I (original single channel 3D image stack)

Output: Final_segmentation (binary image of segmentation result)

Constant Parameters:

     normalization_param = [8000]

     G2_param = [1]

     S3_param = [[1,0.012]]

     min_size = 4

# pre-processing

I_norm = Auto_Contrast(I, normalization_param)

I_smooth = Gaussian_Smoothing_2D_slice_by_slice(I_norm, G2_param)

# apply S3 filter

PreSeg = Spot3D(I_smooth, S3_param)

# watershed to cut falsely merged dots

Mask = Size_Filter(PreSeg, min_size)

Seed = Dilation( Local_maximum(I_norm), structure_element =
       ball_radius_1)

Watershed_map = -1 * Euclidean_distance_transform(Mask)

Watershed_seg = Watershed(Watershed_map, Seed, Mask)

# size filtering

Final_segmentation = Size_Filter(Watershed_seg>0, min_size)
```

### 9. Pseudocode of classic image segmentation workflow for PMP34

```
Input: I (original single channel 3D image stack)

Output: Final_segmentation (binary image of segmentation result)

Constant Parameters:

     normalization_param = [6000]

     G2_param = [1]

     S3_param = [[1,0.03]]

     min_size = 5
```

```
# pre-processing
I_norm = Auto_Contrast(I, normalization_param)
I_smooth = Gaussian_Smoothing_2D_slice_by_slice(I_norm, G2_param)
# apply S3 filter
PreSeg = Spot3D(I_smooth, S3_param)
# watershed to cut falsely merged dots
Mask = Size_Filter(PreSeg, min_size)
Seed = Dilation( Local_maximum(I_norm), structure_element =
       ball_radius_1)
Watershed_map = -1 * Euclidean_distance_transform(Mask)
Watershed_seg = Watershed(Watershed_map, Seed, Mask)
# size filtering
Final_segmentation = Size_Filter(Watershed_seg>0, min_size)
```

### 10. Pseudocode of classic image segmentation workflow for connexin-43

```
Input: I (original single channel 3D image stack)
Output: Final_segmentation (binary image of segmentation result)
Constant Parameters:
     normalization_param = [1, 40]
     G2_param = [1]
     S3_param = [[1,0.031]]
     min_size = 5
# pre-processing
I_norm = Auto_Contrast(I, normalization_param)
I_smooth = Gaussian_Smoothing_2D_slice_by_slice(I_norm, G2_param)
# apply S3 filter
Seg = Spot3D(I_smooth, S3_param)
# size filtering
Final_segmentation = Size_Filter(Seg, min_size)
```

### 11. Pseudocode of classic image segmentation workflow for beta catenin

```
Input: I (original single channel 3D image stack)
Output: Final_segmentation (binary image of segmentation result)
Constant Parameters:
    normalization_param = [4, 27]
    G3_param = [1]
    S2_param = [[1.5,0.01]]
    min_size = 10
# pre-processing
I_norm = Auto_Contrast(I, normalization_param)
I_smooth = Gaussian_Smoothing_3D(I_norm, G3_param)
# apply G2 filter
Seg = Spot2D(I_smooth, S2_param)
# size filtering
Final_segmentation = Size_Filter(Seg, min_size)
```

### 12. Pseudocode of classic image segmentation workflow for tight junction protein ZO1

```
Input: I (original single channel 3D image stack)
Output: Final_segmentation (binary image of segmentation result)
Constant Parameters:
    normalization_param = [3, 17]
    G3_param = [1]
    F3_param = [[1.5,0.2]]
    min_size = 15
# pre-processing
I_norm = Auto_Contrast(I, normalization_param)
I_smooth = Gaussian_Smoothing_3D(I_norm, G3_param)
# apply F3 filter
Seg = Filament3D(I_smooth, F3_param)
```

```
# size filtering
Final_segmentation = Size_Filter(Seg, min_size)
```

**13. Pseudocode of classic image segmentation workflow for beta actin**

```
Input: I (original single channel 3D image stack)
Output: Final_segmentation (binary image of segmentation result)
Constant Parameters:
     normalization_param = [3, 15]
     F3_param = [[2,0.1],[1,0.04]]
     min_size = 15
# pre-processing
I_norm = Auto_Contrast(I, normalization_param)
I_smooth = Edge_Preserving_Smoothing(I_norm)
# apply F3 filter
Seg = Filament3D(I_smooth, F3_param)
# size filtering
Final_segmentation = Size_Filter(Seg, min_size)
```

**14. Pseudocode of classic image segmentation workflow for non-muscle myosin IIB**

```
Input: I (original single channel 3D image stack)
Output: Final_segmentation (binary image of segmentation result)
Constant Parameters:
     normalization_param = [2.5, 17]
     F3_param = [[2,0.2],[1,0.015]]
     min_size = 16
# pre-processing
I_norm = Auto_Contrast(I, normalization_param)
I_smooth = Edge_Preserving_Smoothing(I_norm)
# apply F3 filter
Seg = Filament3D(I_smooth, F3_param)
```

```
# size filtering
Final_segmentation = Size_Filter(Seg, min_size)
```

**15. Pseudocode of classic image segmentation workflow for alpha-actinin-1**

```
Input: I (original single channel 3D image stack)
Output: Final_segmentation (binary image of segmentation result)
Constant Parameters:
    normalization_param = [3, 15]
    F3_param = [[2,0.15], [1,0.05]]
    min_size = 5
# pre-processing
I_norm = Auto_Contrast(I, normalization_param)
I_smooth = Edge_Preserving_Smoothing(I_norm)
# apply F3 filter
Seg = Filament3D(I_smooth, F3_param)
# size filtering
Final_segmentation = Size_Filter(Seg, min_size)
```

**16. Pseudocode of classic image segmentation workflow for alpha tubulin**

```
Input: I (original single channel 3D image stack)
Output: Final_segmentation (binary image of segmentation result)
Constant Parameters:
    normalization_param = [1.5, 8.0]
    F3_param = [[1,0.01]]
    min_size = 20
# pre-processing
I_norm = Auto_Contrast(I, normalization_param)
I_smooth = Edge_Preserving_Smoothing(I_norm)
# apply F3 filter
Seg = Filament3D(I_smooth, F3_param)
```

```
# size filtering
Final_segmentation = Size_Filter(Seg, min_size)
```

**17. Pseudocode of classic image segmentation workflow for troponin I, slow skeletal muscle**

```
Input: I (original single channel 3D image stack)
Output: Final_segmentation (binary image of segmentation result)
Constant Parameters:
    normalization_param = [2, 11]
    F3_param = [[1,0.01]]
    min_size = 15
# pre-processing
I_norm = Auto_Contrast(I, normalization_param)
I_smooth = Edge_Preserving_Smoothing(I_norm)
# apply F3 filter
Seg = Filament3D(I_smooth, F3_param)
# size filtering
Final_segmentation = Size_Filter(Seg, min_size)
```

**18. Pseudocode of classic image segmentation workflow for Titin**

```
Input: I (original single channel 3D image stack)
Output: Final_segmentation (binary image of segmentation result)
Constant Parameters:
    normalization_param = [8, 15.5]
    F3_param = [[1,0.02]]
    min_size = 15
# pre-processing
I_norm = Auto_Contrast(I, normalization_param)
I_smooth = Edge_Preserving_Smoothing(I_norm)
# apply F3 filter
Seg = Filament3D(I_smooth, F3_param)
```

```
# size filtering
Final_segmentation = Size_Filter(Seg, min_size)
```

**19. Pseudocode of classic image segmentation workflow for lamin B1 (interphase-specific)**

```
Input: I (original single channel 3D image stack)
Output: Final_segmentation (binary image of segmentation result)
Constant Parameters:
    normalization_param = [4000]
    G3_param = [1]
    mid_stack_method = 'intensity'
    F2_param = [[1,0.01], [2,0.01], [3,0.01]]
    seed_param = [400, 40000]
# pre-processing
I_norm = Auto_Contrast(I, normalization_param)
I_smooth = Gaussian_Smoothing_3D(I_norm, G3_param)
# get the middle slice of the stack
Mid_z = GetMidStack(I_smooth, mid_stack_method)
# apply F2 filter
Seg_mid_z = Filament2D(I_smooth at Mid_z, F2_param)
# apply watershed to get shells
Seed_img = Logical_XOR(Seg_mid_z, Hole_Filling(Seg_mid_z,
           seed_param))
Seed = Cetroids of connected_component(Seed_img)
Final_segmentation = Watershed(I_norm, Seed)>0
```

**20. Pseudocode of classic image segmentation workflow for lamin B1 (mitosis-specific)**

```
Input: I (original single channel 3D image stack)
Output: Final_segmentation (binary image of segmentation result)
Constant Parameters:
    normalization_param = [4000]
```

```
    G3_param = [1]
    F2_param = [[0.5,0.01]]
    min_size = 20
# pre-processing
I_norm = Auto_Contrast(I, normalization_param)
I_smooth = Gaussian_Smoothing_3D(I_norm, G3_param)
# apply F2 filter
Seg = Filament2D(I_smooth, F3_param)
# size filtering
Final_segmentation = Size_Filter(Seg, min_size)
```
